# Signal sequences encode information for protein folding in the endoplasmic reticulum

**DOI:** 10.1101/2020.06.04.133884

**Authors:** Sha Sun, Xia Li, Malaiyalam Mariappan

## Abstract

One-third of newly synthesized proteins in mammals are translocated into the endoplasmic reticulum (ER) through the Sec61 translocon. It remains unclear how protein translocation coordinates with the chaperone availability to promote protein folding in the ER. We find that signal sequences cause a translocation pause at the Sec61 translocon until nascent chains engage with luminal chaperones for efficient translocation and folding in the ER. Using a substrate-trapping proteomic approach, we identify that nascent proteins with marginally hydrophobic signal sequences accumulate on the cytosolic side of the Sec61 translocon, which recruits BiP chaperone through Sec63 to bind onto nascent chains. Surprisingly, BiP binding not only releases translocationally paused nascent chains into the ER lumen but also ensures protein folding. Increasing hydrophobicity of signal sequences bypasses Sec63/BiP-dependent protein translocation but translocated nascent chains misfold and aggregate under conditions of limited BiP availability in the ER. Thus, signal sequence-dependent protein folding explains why signal sequences are diverse and use multiple protein translocation pathways.

## Introduction

About 30% of newly synthesized proteins from the cytosolic ribosomes are delivered to the endoplasmic reticulum (ER) by the co-translational protein targeting pathway (Juszkiewicz and Hegde, 2018). These nascent polypeptides typically contain either cleavable N-terminal signal sequences (SSs) or non-cleavable first transmembrane domains (TMDs). As SSs emerge from cytosolic ribosomes, they are co-translationally recognized and captured by the signal recognition particle (SRP) (Shan and Walter, 2005). The SRP-bound ribosome-associated nascent chains (RNCs) are then delivered to the ER membrane by an interaction with the SRP receptor (Jiang et al., 2008). The SS is then transferred from the SRP to the heterotrimeric Sec61 translocon complex (Rapoport, 2007). The SS is again recognized, and bound, this time, by the Sec61 translocon, allowing the translocation of the following nascent polypeptide across the ER membrane with concomitant cleavage of a signal peptide or integration of a TMD into the lipid bilayer.

SSs are diverse and significantly differ in length, hydrophobicity, charge, and flanking mature domain, but all of them contain a core hydrophobic region of at least six non-hydrophilic residues. Earlier studies have shown that SRP can efficiently recognize a broad range of hydrophobic SSs of RNCs in the cytosol and target them to the ER of mammalian cells (Hegde and Kang, 2008; Kim et al., 2002; Voorhees and Hegde, 2015). However, the interaction between the Sec61 translocon and SSs greatly varies. Surprisingly, only a minority of signal sequences, such as the signal sequence from prolactin, tightly engages with the Sec61 translocon. This results in a looped orientation in which the N-terminus of the SS is positioned at the cytosolic face of the translocon, and the following the hydrophobic portion of the SS becomes intercalated into the lateral gate of the Sec61 translocon (Jungnickel and Rapoport, 1995; Kim et al., 2002; Voorhees and Hegde, 2016a). Hence, these SSs mediate efficient protein translocation through the Sec61 translocon. Surprisingly, most SSs inefficiently interact with the Sec61 translon, leading to a nonlooped orientation that forces the mature domain of a nascent chain extruded into the cytosol (Hegde and Kang, 2008). These signal sequences require accessory factors including TRAM, TRAP, Sec63/Sec63, and luminal chaperones to improve their interaction with the Sec61 translocon and thereby promote the translocation of their mature domains into the ER (Brodsky et al., 1995; Fons et al., 2003; Nguyen et al., 2018; Schorr et al., 2020b; Voigt et al., 1996). If the role of signal sequences is merely to mediate targeting and translocation into the ER, it is difficult to rationalize why signal sequences are evolved to use different accessory factors for their efficient translocation into the ER. One important function of SSs is that protein translocation into the ER can be selectively attenuated in a signal sequence-dependent manner during ER stress to reduce the ER folding burden (Kang et al., 2006). In some cases, the SS can regulate the timing of signal peptide cleavage, thereby enhancing protein maturation (Rehm et al., 2001; Rutkowski et al., 2003; Snapp et al., 2017). However, the factors and mechanisms involved in the SS-dependent protein maturation are less understood.

Similar to SSs, TMDs of polytopic membrane proteins are recognized by the Sec61 translocon and laterally partition into the lipid bilayer using the lateral gate of the translocon. Again, TMD hydrophobicity plays a crucial role in partitioning from the lateral gate of the translocon to the hydrophobic lipid bilayer. Yet, TMDs of membrane proteins greatly vary in their hydrophobicity. It is estimated that ~25% of TMDs of polytopic membrane proteins are marginally hydrophobic, meaning they are not suitable for insertion into the membrane by themselves in isolation (Elofsson and von Heijne, 2007; White and von Heijne, 2008). Recent studies have shown that marginally hydrophobic can transiently stall in the interior pore of the Sec61 translocon (Kida et al., 2016; Kida and Sakaguchi, 2018). It is unclear how and which factors assist in clearing the marginally hydrophobic TMD that is transiently retained at the Sec61 transocon during the co-translational membrane protein insertion. In some cases, less hydrophobic TMDs are completely translocated into the ER lumen (Feige and Hendershot, 2013; Ojemalm et al., 2012; Skach et al., 1994; Sun and Mariappan, 2020). They can then either be retrieved by interactions with the membrane embedded TMDs or degraded by ER-associated quality control pathways (Buck et al., 2017).

Upon translocation into the ER lumen, nascent chains are folded with the assistance of molecular chaperones and folding enzymes (Braakman and Hebert, 2013; Fewell et al., 2001; Helenius et al., 1992; Ma and Hendershot, 2004). Among the broad spectrum of chaperones in the ER, the member of the Hsp70 family chaperone BiP ATPase plays a central role in shielding exposed hydrophobic residues in nascent proteins and thereby preventing protein misfolding and aggregation. BiP also assists in protein folding by cycling between high-affinity and low-affinity substrate binding through ATP-dependent conformational changes that are regulated by co-chaperones and nucleotide exchange factors in the ER lumen (Pobre et al., 2019).

Considering the essential roles of BiP in preventing protein misfolding and promoting folding, it is less understood how the translocating nascent chains are accurately matched to the availability of active BiP molecules in the ER lumen. Studies of cytosolic proteins have shown that the ribosome-associated complex (RAC) recruits Hsp70 to assist the folding of aggregation-prone nascent polypeptides (Doring et al., 2017; Willmund et al., 2013). It is unclear whether a similar mechanism exists in the ER since the ribosome is separated from the translocating nascent chain by the ER membrane bilayer. This physical barrier suggests that the Sec61 translocon and its associated factors (Gemmer and Forster, 2020) might recruit the luminal the Hsp70 chaperone BiP to the aggregation-prone nascent polypeptides for their proper folding and maturation in the ER.

In this study, we used a series of substrates that vary in TMD hydrophobicity to demonstrate that the Sec61 translocon-associated membrane protein Sec63 is responsible for releasing marginally hydrophobic (or weak) TMDs that are transiently retained at the Sec61 translocon by recruiting and activating BiP ATPase. We developed a substrate-trapping proteomic approach and identified endogenous secretory and membrane proteins that are transiently retained at the Sec61 translocon in the absence of Sec63 activity. We demonstrate that the nascent chain with a marginally hydrophobic (or weak) SS is co-translationally recruited to the ER membrane, but the initial translocation of ~160 amino acids is paused and is accumulated on the cytosolic side of the Sec61 translocon. Sec63 releases the translocation pause through recruiting and activating BiP ATPase to bind and facilitate the nascent chain translocation into the ER. Surprisingly, the BiP-mediated co-translational translocation also guarantees subsequent protein folding in the ER. Increasing the hydrophobicity of a weak SS can bypass the Sec63/BiP-dependent translocation, but the translocated nascent chain is at a high risk of misfolding and aggregating under conditions of limited BiP availability in the ER. Thus, our studies suggest that SSs have evolved to recruit appropriate chaperones through modulating translocon-associated factors to meet the unique challenges of folding secretory and membrane proteins in the ER.

## Results

### The Sec63/BiP complex clears translocons transiently retained with marginally hydrophobic sequences

We first investigated how the Sec61 translocon responds to marginally hydrophobic sequences in proteins during their translocation into the ER. Earlier studies have shown such sequences are neither hydrophobic enough to be inserted into the lipid bilayer nor hydrophilic enough to be translocated into the ER lumen, leading to their transient retention at the Sec61 translocon (Kida et al., 2016; Kida and Sakaguchi, 2018). To investigate the mechanism by which the transiently retained nascent chain at the Sec61 translocon is cleared, we used our model C-terminally Venus-tagged substrates derived from WRB, a subunit of the tail-anchored membrane protein insertase (Fig. S1A) (Yamamoto and Sakisaka, 2012). We have recently shown that the C-terminal TMD of WRB-Venus is translocated into the ER lumen because of a positively charged lysine residue (Fig. S1A) (Sun and Mariappan, 2020). The translocation of the C-TMD of WRB into the ER lumen can be further increased by either incorporating more charged residues or reduced by adding more hydrophobic residues (Sun and Mariappan, 2020) (Fig. S1B).

Consistent with our recent findings, substrates with marginally hydrophobic C-terminal TMDs (K, D, RRK, and DDRRK), provisionally termed weak TMDs, were poorly inserted into the ER membrane and were translocated into the ER lumen as evidenced by the glycosylation (Fig. 1A and 1B and Fig. S1B). By contrast, the substrate with sufficient hydrophobicity (L) was efficiently inserted into the ER membrane as shown by the mostly non-glycosylated form (Fig. 1A and 1B).

**Figure 1.**
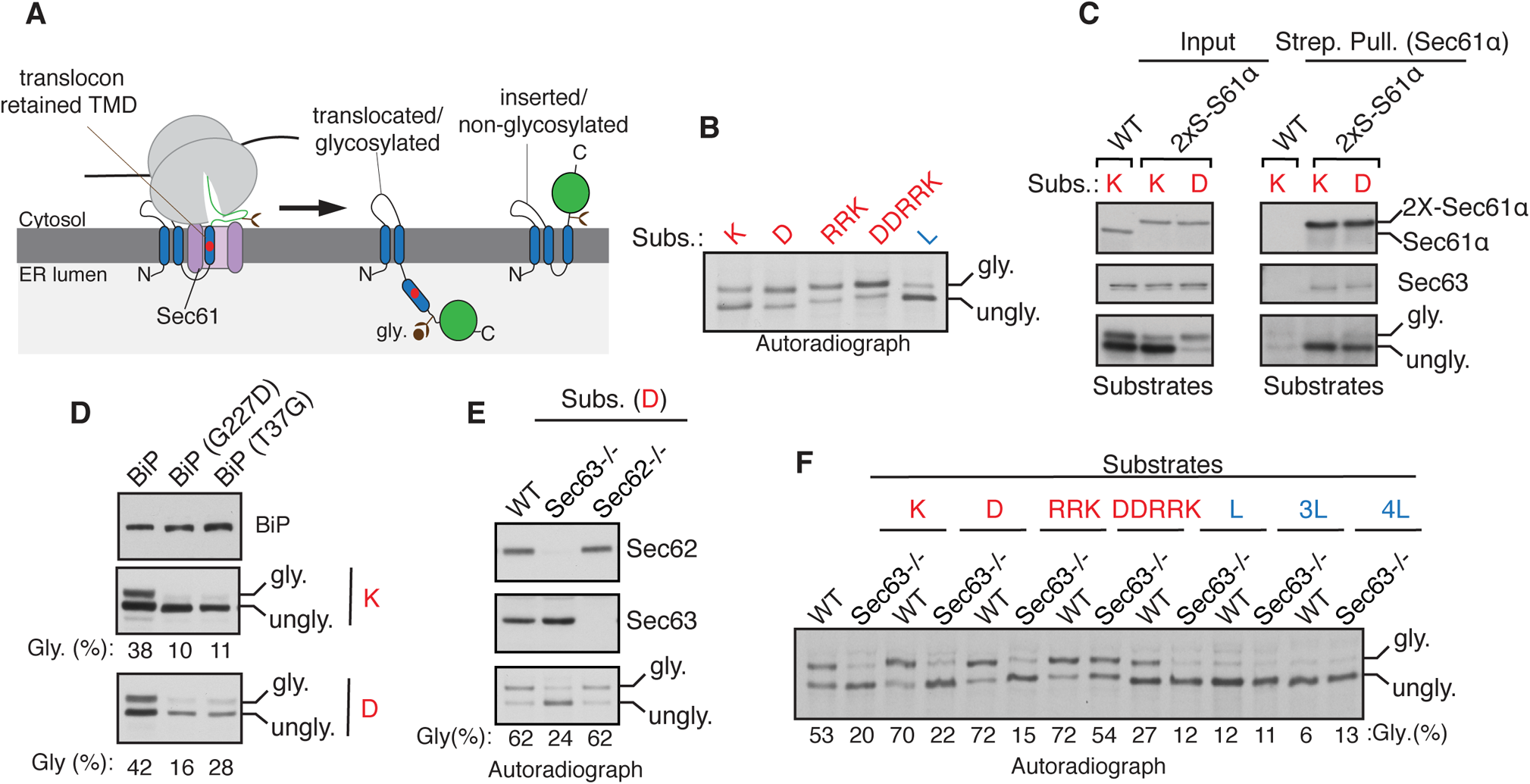
Sec63 is required for releasing marginally hydrophobic TMDs retained at the Sec61 translocon. (A) A diagram illustrating the translocon retained with the marginally hydrophobic C-terminal TMD of WRB-Venus, which is then translocated into the ER lumen and is glycosylated. The C-TMD with sufficient hydrophobicity is properly inserted into the ER membrane and thereby is not glycosylated. The red circle in the C-TMD indicates a charged residue. (B) Cells expressing C-TMDs of substrates bearing indicated charged residues (shown in red color) or hydrophobic residue (indicated in blue color) were radiolabeled and analyzed by SDS-PAGE and autoradiography after immunoprecipitation of substrates with anti-GFP antibodies. (C) The indicated substrates were expressed in chromosomally 2x Strep-tagged Sec61α cells, lysed, and pulled down with Strep-Tactin beads and analyzed by immunoblotting for the indicated antigens. The substrates were immunoblotted with anti-GFP antibodies. (D) Cells co-expressing the indicated versions of BiP and the substrate were analyzed by immunoblotting with anti-BiP antibodies or anti-GFP antibodies for substrates. The percent of glycosylation is indicated below each panel. (E) WT HEK293, Sec62−/−, or Sec63−/− cells expressing the indicated substrates were radiolabeled and analyzed as in (B). Immunoblots show the knockout of either Sec62 or Sec63. (F) WT HEK293 or Sec63−/− cells expressing the indicated substrates were immunoprecipitated with anti-GFP antibodies and analyzed by autoradiography.

We hypothesized that weak TMD substrates may be transiently retained at the Sec61 translocon before their translocation into the ER lumen. To test this, we precipitated the chromosomally 2xStrep-tagged Sec61α from cells expressing weak TMD substrates and found that the unglycosylated form was highly enriched with the Sec61 translocon (Fig. 1C). However, the glycosylated form was not significantly associated with the Sec61 translocon since it was already translocated into the ER lumen and likely moved away from the translocon. Next, we asked how the transiently retained substrates are cleared from Sec61 translocons. We hypothesized that the abundant luminal chaperone BiP ATPase might promote the translocation of transiently retained substrates from Sec61 translocons. Indeed, the transient expression of BiP mutants (G227D and T37G), which are defective in ATP-induced release of substrates (Wei et al., 1995), in cells inhibited the translocation of weak TMDs into the ER lumen (Fig. 1D).

The BiP involvement in this process raises the question of how BiP is locally recruited to the Sec61 translocon to release weak TMDs into the ER lumen. The translocon-associated membrane protein Sec63 is known to recruit and activate BiP ATPase in order to facilitate post-translational protein translocation into the ER lumen in yeast (Deshaies et al., 1991; Matlack et al., 1999; Meyer et al., 2000). To test if Sec63 is required in this process, we used knockout cells of either Sec63 or its interacting protein Sec62 generated by CRISPR/Cas9. Although both Sec62 and Sec63 are required for BiP-mediated post-translational translocation into the ER, Sec63 was sufficient to translocate the translocon-retained weak TMD into the ER lumen (Fig. 1E). To determine biochemical features of TMDs that depend on Sec63 activity, we tested the translocation of TMDs with varying hydrophobicity in Sec63−/− cells (Fig. 1F and S1B). The translocation of substrates with weak TMDs (K, D, and RRK) was significantly impaired as demonstrated by reduced glycosylated forms in Sec63−/− cells relative to that of wild-type (WT) cells. By contrast, strong TMDs (L, 3L and 4L) were mostly inserted into the membrane as evidenced by the appearance of predominantly non-glycosylated forms in both WT and Sec63−/− cells (Fig. 1F). Of note, the extremely charged TMD (DDRRK) was only partially dependent on Sec63 (Fig. 1F). These results suggest that a weak TMD transiently retains at the Sec61 translocon and thereby relies on Sec63 activity for its translocation into the lumen, whereas Sec63 activity is dispensable if the TMD is either sufficiently hydrophobic or it is changed to a nearly hydrophilic sequence. Consistent with our data from cells, the translocation of a charged

TMD into the ER lumen depended on Sec63 activity in an in vitro assay performed using microsomes derived from WT or Sec63−/− cells (Fig. S1C). Surprisingly, Sec63 was dispensable for translocation of a weak TMD with a small C-terminal tail containing 26 amino acids (Fig. S1D). This result indicated that the C-terminal tail length after the TMD is crucial for its Sec63-dependent translocation into the ER lumen. Truncation studies further revealed that the translocation of the C-TMD with a tail longer than 100 amino acids was dependent on Sec63 activity in cells (Fig. S1D). Taken together, these data suggest that Sec63 is required for clearing the Sec61 translocon transiently retained with weak TMDs containing the C-terminal tails longer than 100 amino acids.

### A substrate-trapping approach identifies the transiently retained endogenous substrates at the Sec61 translocon

To determine the role of Sec63 in clearing the translocon of retained endogenous substrates, we developed a substrate trapping strategy. We predicted that a Sec63 J-domain mutant could trap the Sec61 translocon-retained with nascent chains because the mutant J-domain cannot recruit and activate BiP ATPase. To examine this, we stably complemented either WT Sec63, Sec63 J-mutant (HPD/AAA), or the translocon interaction defective Sec63 mutant (Δ230-300) into Sec63−/− cells (Fig. S2A). Using our model substrates, we first tested whether Sec63 J-domain or its interaction with the translocon is essential for clearing the transiently retained substrates at the Sec61 translocon. WT Sec63 complemented cells cleared translocons retained with weak TMDs by releasing them into the ER lumen as illustrated by the glycosylated form of substrates (Fig. 2A and S2B). By contrast, Sec63 J-mutant and the Sec61 interaction defective mutant (Δ230-300) were not able to clear the translocon-retained substrates as evidenced by the primarily unglycosylated form of substrates (Fig. 2A and S2B). Using these stable cell lines, we next performed metabolic labeling and immunoprecipitation to determine if Sec63 J-mutant could trap translocons-retained with endogenous substrates, and if the trapping depends on its interaction with the translocon. We found that all three subunits of the newly synthesized Sec61 translocon, α, β, and γ were associated with both WT Sec63 and J-mutant but were not detected from either Sec63−/− cells or the Sec61 interaction defective mutant (Δ230-300) expressing cells (Fig. 2B). Quantification of bands further revealed that newly synthesized radiolabeled proteins were significantly enriched with Sec63 J-mutant relative to WT Sec63 or Sec63 (Δ230-300) (Fig. 2B). These findings support our conclusion that both Sec63 J-domain and its interaction with the translocon are important for clearing translocons transiently retained with nascent polypeptides in cells.

**Figure 2.**
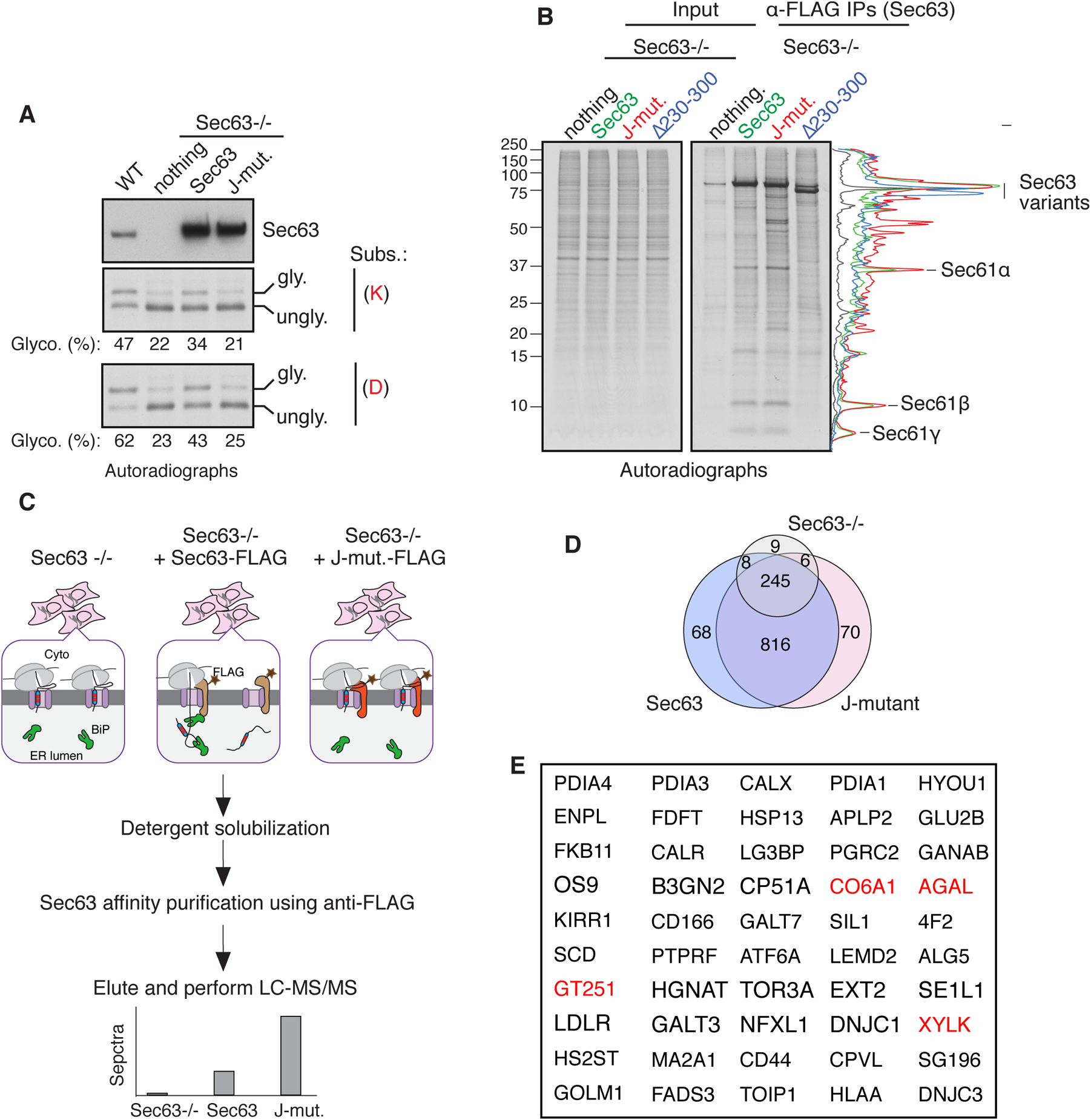
Identification of translocon-retained endogenous substrates of Sec63. (A) WT HEK293 or Sec63−/− cells stably expressing the indicated Sec63 constructs were transfected with the C-TMD of WRB substrate carrying either lysine (K) or aspartic acid (D). The transfected cells were radiolabeled and analyzed by autoradiography after immunoprecipitation with anti-GFP antibodies for substrates. The immunoblot shows the expression of Sec63 from the indicated cell lines. (B) The indicated Sec63−/− complemented cell lines were radiolabeled and analyzed by autoradiography after immunoprecipitation with anti-FLAG beads. The traces are densitometry profiles of lanes and shown at the right side of the autoradiograph, illustrating a selective enrichment of signals with the Sec63 J-mutant (HPD/AAA). (C) A scheme describing affinity purification and identification of translocon-retained substrates by mass spectrometry. (D) Venn diagram of protein distribution among the three groups. The diagram was built from mass spectrometry data from Table S1. (E) The top 50 secretory and membrane proteins from Table S2 that were enriched in Sec63 J-mutant. The red color labeled proteins are found also in the study by Schorr et al., 2020.

We next performed a large-scale affinity purification of Sec63 as depicted in Fig. 2C and subjected immunoprecipitants to mass spectrometry to identify translocon-associated endogenous clients. Our analysis showed many proteins associated with both WT Sec63 and Sec63 J-mutant relative to control Sec63−/− cells (Fig. 2D and Table S1). Consistent with our prediction, about 117 secretory and membrane proteins were at least two-fold enriched in Sec63 J-mutant relative to WT Sec63 and five-fold enriched than Sec63−/− cells (Fig. 2E and Table S2). Of the 117 proteins, about 65% of clients were non-resident ER proteins, indicating that these clients interacted with Sec63 J-mutant during their retention at the Sec61 translocon in the ER. Interestingly, about one-third of these candidate substrates have been previously implicated in human genetic disorders caused by mutations in corresponding genes (Table S2). Many of our Sec63 clients were also found in the independent proteomic study performed using Sec63 depleted cells (Fig. 2E and Table S2) (Schorr et al., 2020a). We found about 86 secretory and membrane proteins were enriched with WT Sec63, but about 60% of them are ER-resident proteins, suggesting that many of these proteins are likely interacting proteins of Sec63 in the ER (Table S3). Thus, our substrate-trapping proteomic approach identified potential endogenous secretory and membrane proteins that are transiently retained at the Sec61 translocon in the absence of Sec63 activity.

### Proteins containing weak signal sequences depend on Sec63 for their translocation into the ER lumen

To validate our Sec63’s substrates, we constructed C-terminally FLAG-tagged expression plasmids for two substrates, AGAL and CD44 (Fig. S3A and S3B). The *AGAL* gene encodes for a lysosomal enzyme called α-galactosidase A, the activity of which is altered by genetic mutations in Fabry disease (Germain, 2010). The *CD44* gene encodes a plasma membrane-localized single spanning membrane protein with an N-terminal SS. We first examined if these substrates depended on Sec63 to translocate into the ER. Immunoblotting of AGAL from transiently expressing cells revealed that AGAL was fully glycosylated (Fig. 3A). AGAL glycosylation was verified by digestion with Endo H, which removes high mannose N-linked glycans from glycoproteins. AGAL glycosylation was markedly reduced in Sec63−/− cells relative to WT cells (Fig. 3A). Surprisingly, Endo H digestion revealed that unglycosylated AGAL in Sec63−/− cells migrated slower than Endo H-deglycosylated AGAL (Fig. 3A and Fig. S3C), which suggests that the unglycosylated AGAL in Sec63−/− cells still contains the uncleaved SS. The unglycosylated AGAL carrying the SS was mislocalized to the cytosol of Sec63−/− cells, as it was accessible to proteinase K (PK) added to the semipermeabilized cells, but the translocated/glycosylated form was mostly protected from PK digestion (Fig. S3D). Expression of CD44 in WT cells showed two glycosylated bands. The faster migrating core glycosylated band was sensitive to Endo H but the slower migrating complex type of N-glycan was less sensitive to Endo H, indicating that it is localized to post-ER compartments (Fig. 3A). Similar to AGAL, the translocation of CD44 was significantly reduced as it displayed mostly the unglycosylated form in Sec63−/− cells. We then asked if the translocation defects associated with AGAL and CD44 could be rescued by WT Sec63. Indeed, both AGAL and CD44 were translocated at similar levels to that of WT cells when Sec63−/− cells were complemented with WT Sec63 (Fig. 3B). By contrast, Sec63 J-mutant (HPD/AAA) complemented cells showed almost no translocation of AGAL and CD44 relative to WT Sec63 complemented cells and Sec63−/− cells. (Fig. 3B). This result implies that the activation of BiP ATPase by the J-domain of Sec63 is critical for the translocation of these proteins. Since there is no glycosylated form of AGAL in the J-mutant lane, this result also suggests that Sec63 J-mutant completely inhibits Sec63 independent translocation in Sec63−/− cells (Fig. 3B), presumably by blocking spontaneous BiP binding to the translocating nascent chains. Since Sec63 functions along with Sec62 in post-translational protein translocation, we wondered whether Sec62 is also required for translocation of AGAL and CD44. While the knockout of Sec62 had a significant effect on the translocation of CD44 as mirrored by less glycosylated forms, it had a subtle effect on the translocation of AGAL, suggesting that Sec62 plays a substrate-dependent role in promoting translocation of secretory proteins (Fig. 3C).

**Figure 3.**
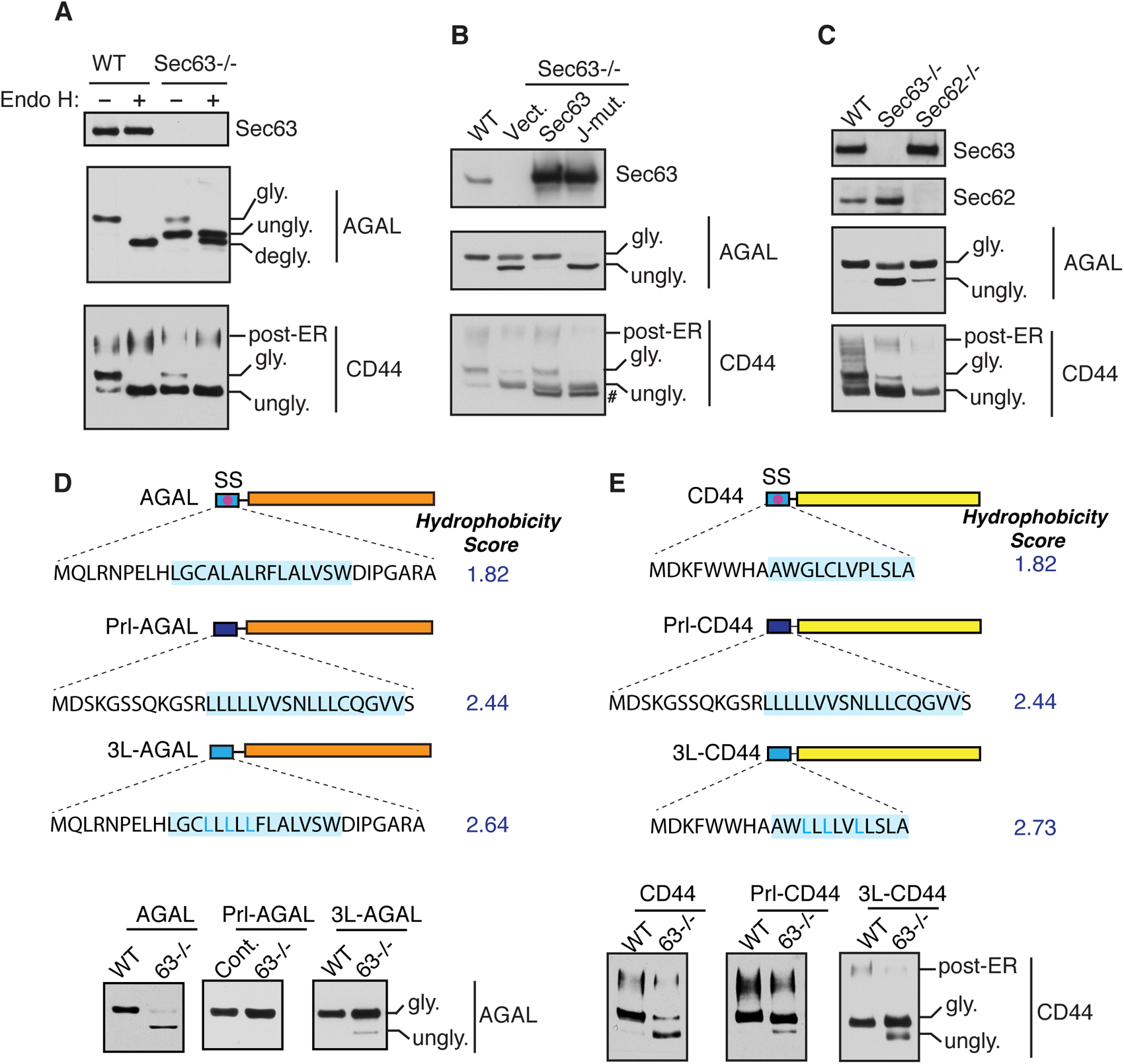
Translocation of proteins with less hydrophobic signal sequences depend on Sec63. (A) WT HEK293 or Sec63−/− cells were transfected with AGAL-FLAG or CD44-FLAG. Cell lysates were either untreated or treated with Endo H and analyzed by immunoblotting with an anti-FLAG antibody for substrates and anti-Sec63 antibodies. “gly” indicates glycosylated forms. “ungly” denotes non-translocated/unglycosylated forms with uncleaved SSs. “degly” indicates Endo H digested de-glycosylated forms. Post-ER form of CD44 represents the Endo H resistant glycosylated form. (B) WT HEK293 or Sec63−/− cells stably expressing the indicated constructs were transfected with AGAL-FLAG or CD44-FLAG. Cell lysates were directly analyzed by immunoblotting as in (A). # denotes FLAG-tagged Sec63 proteins. (C) WT HEK293, Sec63−/−, or Sec62−/− cells were transfected with plasmids expressing AGAL-FLAG or CD44-FLAG, and the cell lysates were directly analyzed as in A. (D) Top: Cartoons depicting the signal sequence (SS) of AGAL is either replaced with the SS of prolactin (Prl-AGAL) or mutated to generate 3L-AGAL. Blue color shade denotes the hydrophobic region (h-region) of the signal sequence. Hydrophobicity scores of h-regions were analyzed by grand average hydropathy (GRAVY). Bottom: The indicated cell lines were transfected with the mentioned constructs and analyzed by immunoblotting with an anti-FLAG antibody. (E) Top: A cartoon depicting the SS of CD44 that is either replaced with the SS of prolactin (Prl-CD44) or mutated to generate 3L-CD44. Hydrophobicity scores were analyzed as in (D). The indicated cell lines were transfected and analyzed as in D.

Since our data above showed that weak TMDs relied on Sec63 activity (Fig. 1), we wanted to determine if our new Sec63 substrates AGAL and CD44 also contain weak SSs. We therefore swapped the SS of AGAL or CD44 with the strong SS from secretory hormone prolactin (prl) and tested its Sec63 dependency by expressing it in Sec63−/− cells (Fig. 3D). Prl-AGAL bypassed the Sec63 requirement as demonstrated by a fully glycosylated form of AGAL in both WT and Sec63−/− cells, whereas AGAL with its endogenous SS poorly glycosylated in Sec63−/− cells (Fig. 3D). Interestingly, some of the hydrophilic residues within the core hydrophobic region of AGAL SS are conserved in mammals, suggesting an evolutionary pressure to maintain Sec63 dependency (Fig. S4A). Indeed, the hydrophobicity is the key determinant for Sec63 dependency since the translocation could occur independent of Sec63 activity for 3L-AGAL in which less hydrophobic amino acids of AGAL SS were replaced with hydrophobic leucine amino acids (Fig. 3D). We also obtained a similar result for CD44 where Prl-CD44 and 3L-CD44 mostly translocated independent of Sec63 activity compared to WT CD44 (Fig. 3E and Fig. S4B). Lastly, replacing prolactin SS with less hydrophobic AGAL SS led to Sec63 dependent translocation of prolactin into the ER, suggesting that the weak SS of AGAL is sufficient to render Sec63 dependency for a protein (Fig. S4C). Collectively, our results show that weak SSs rely on Sec63/BiP for efficient translocation of their nascent chains into the ER lumen.

### Sec63 co-translationally releases the translocation pause caused by a marginally hydrophobic signal sequence

Previous studies have established that Sec63 is predominantly involved in post-translational translocation of substrates into the ER lumen by recruiting and activating BiP ATPase through its J-domain (Corsi and Schekman, 1997; Matlack et al., 1999). Studies have shown that Sec63 is also involved for the co-translational protein translocation of select substrates (Brodsky et al., 1995; Conti et al., 2015; Jung et al., 2014; Lang et al., 2012; Young et al., 2001), but its precise role in this pathway is ill-defined. The recent structural studies of the Sec61/Sec62/Sec63 complex from yeast suggest that both Sec63 and ribosome share the same binding site in the Sec61 translocon (Itskanov and Park, 2019; Wu et al., 2019). This means both the ribosome and Sec63 cannot bind to the Sec61 translocon at the same time, questioning the role of Sec63 in the co-translational translocation process. To determine whether our novel Sec63’s substrates are co- or post-translationally translocated into the ER, we performed in vitro translation of AGAL transcripts using the rabbit reticulocyte (RRL) translation system including [^35^S] methionine in either the presence or absence of ER-derived rough microsomes. AGAL was digested by proteinase K (PK) without microsomes, but it was translocated and glycosylated in the presence of microsomes and thereby insensitive to protease digestion (Fig. 4A, and 4B). In sharp contrast, AGAL was completely digested by PK when microsomes were post-translationally incubated with the supernatant of the translation reaction in which ribosomes were removed by centrifugation (Fig. 4A and 4B). To directly detect the co-translational translocation of AGAL, we translated AGAL transcripts lacking the stop codon in vitro in either the absence or presence of microsomes. The absence of stop codon left the nascent polypeptide attached to the ribosome by a tRNA peptidyl linkage (Fig. 4C). AGAL translated in the absence of microsomes was completely digested by PK. By contrast, AGAL or Prl-AGAL translated in the presence microsomes was translocated into the ER lumen as demonstrated by protease resistant glycosylation (Fig. 4C). We noticed that glycosylation of AGAL into microsomes was slightly less efficient than that of Prl-AGAL. The fact that the nascent chain of AGAL while covalently attached to the ribosome could be translocated and glycosylated supports the conclusion that AGAL is co-translationally translocated into the ER.

**Figure 4.**
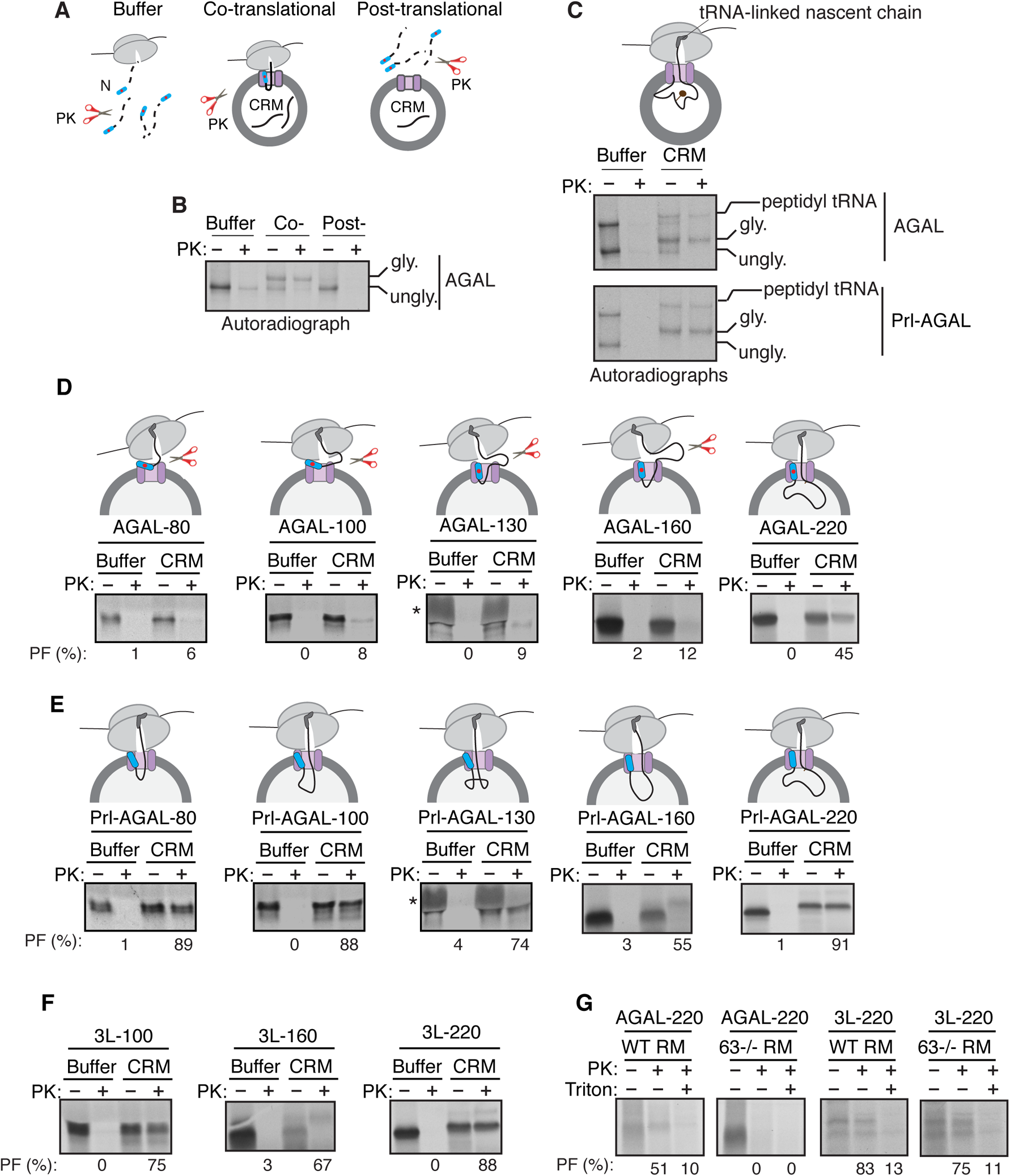
Sec63 substrates are co-translationally translocated with an initial pause. (A) Scheme depicting the protocol for proteinase K (PK) treatment for the co- or post-translational protein translocation assay. (B) For the co-translational protein translocation, transcripts encoding AGAL-FLAG were *in vitro* translated in the rabbit reticulocyte lysate (RRL) based translation system including either buffer or canine rough microsomes (CRM). After translation, samples were digested with PK and analyzed by autoradiography after immunoprecipitation with anti-FLAG antibody. For the post-translational translocation, transcripts encoding AGAL-FLAG were translated and centrifuged to remove ribosomes. The supernatant was incubated with CRM and digested with PK and analyzed after immunoprecipitation with anti-FLAG antibody. (C) AGAL or Prl-AGAL transcripts lacking stop codon were *in vitro* translated in RRL including either buffer or CRM. The reactions were treated with PK and analyzed as in (B). Note that transcripts lacking stop codon produce nascent chains that remain attached to ribosomes through a covalent peptidyl-tRNA bond. (D). The ribosome-associated nascent chains (RNCs) of the indicated lengths of AGAL were produced in RRL including either buffer or CRM, digested with PK, treated with RNase A to remove tRNA and analyzed by autoradiography. The percentage of protease protected fragments (PF) is shown below each panel. The star symbol indicates a distortion of the AGAL-130 nascent chain caused by the co-migration with abundant hemoglobin from RRL. (E and F) RNCs of the indicated lengths of Prl-AGAL or 3L-AGAL were produced and analyzed as in D. (G) Transcripts of AGAL-220 or 3L-AGAL-220 were translated in the presence of either rough microsomes (RM) derived from WT HEK293 cells or Sec63−/− cells and digested with PK before analyzing by autoradiography. Note that protease protected fragments were mostly disappeared in samples including 1% Triton X-100.

The data above show that Sec63-dependent substrate (AGAL) and Sec63-independent substrate (Prl-AGAL) are co-translationally translocated into the ER lumen. We therefore wanted to determine at which step during the biosynthesis of AGAL, Sec63 would assist the translocation of AGAL. To address this, we in vitro translated transcripts lacking the stop codon encoding defined chain lengths of AGAL in either the presence or absence of microsomes (Fig. 4D). Translation of transcripts without stop codon yielded translationally arrested radiolabeled ribosome-associated nascent chains (RNCs), representing physiological translocation intermediates. RNCs of AGAL-80, -100, -130, and 160 amino acids in length were digested by PK, as minimal protease protected fragments were observed (Fig. 4D). However, the RNC of AGAL-220 was protected from PK digestion. These results suggest that the AGAL nascent chain of ~160 amino acids is translocationally paused and accumulated to the cytosolic side of the Sec61 translocon and thus accessible to PK digestion. The translocation pause was released when the nascent chain reaches the length of 220 amino acids, as shown by the appearance of PK protected nascent chains. Conversely, Prl-AGAL, which is not dependent on Sec63, showed protease protected fragments with all different lengths of nascent chains, illustrating that the successful translocation is initiated right after the SS engages the Sec61 translocon (Fig. 4E).

The protease protection experiment with 3L-AGAL nascent chains showed a similar result to that of Prl-AGAL (Fig. 4F), which demonstrates that hydrophobicity of the SS determines the efficient translocation of nascent chains into the ER. The translocation defects associated with AGAL nascent chains below 220 residues were not caused by their inability to target to the ER membrane since both AGAL-100 and AGAL-220 nascent chains were recruited to microsomes with a comparable efficiency to nascent chains of 3L-AGAL (Fig. S5). We next asked whether Sec63 is required for assisting co-translational translocation of AGAL nascent chain into the ER in vitro. Indeed, nascent chains of AGAL-220 were protease sensitive when incubated with microsomes derived from Sec63−/− cells compared to microsomes derived from WT cells (Fig. 4G). In contrast, 3L-AGAL showed protease protected fragments for both WT and Sec63−/− microsomes since its translocation into the ER was independent of Sec63. Collectively, these results suggest that Sec63 substrates are co-translationally translocated into the ER lumen with an initial translocation pause due to their weak SSs. Our data also suggest that Sec63 can release the translocationally paused nascent chain into the ER even in the presence of translating ribosome.

### Sec63 mediates BiP binding to translocating nascent chains in the ER

Next, we asked why weak SSs are frequently found in secretory proteins despite their inefficiency in mediating translocation. We hypothesized that SSs may direct the recruitment of luminal chaperones to bind onto nascent chains through modulating the Sec61 translocon associated factors. Accordingly, we predicted that the weak SS of AGAL nascent chain would preferentially bind to BiP chaperone since its translocation depends on the J-domain of Sec63. To test this, we introduced a C-terminal 3xFLAG-tag on endogenous genes encoding BiP using CRISPR/Cas9 because commercial antibodies against BiP could not efficiently pulldown the endogenous BiP. The C-terminal 3xFLAG introduced downstream of the KDEL retention signal of BiP did neither significantly change BiP expression levels nor its localization to the ER (Fig. 5A, B). To determine if the newly synthesized AGAL nascent chain is a substrate of BiP, BiP-3xFLAG HEK293 cells expressing AGAL were radiolabeled with ^35^S methionine/cysteine for 2 min, chased, and immunoprecipitated with an anti-FLAG antibody. Consistent with our prediction, AGAL nascent chains were co-immunoprecipitated with BiP. The interaction reduced with the chase time because matured AGAL presumably left the ER (Fig. 5C). BiP binding to AGAL and other endogenous substrates was sensitive to ATP since the incubation of cell lysates with ATP prior to immunoprecipitation completely abolished BiP binding to all nascent chains including AGAL (Fig. 5D).

**Figure 5.**
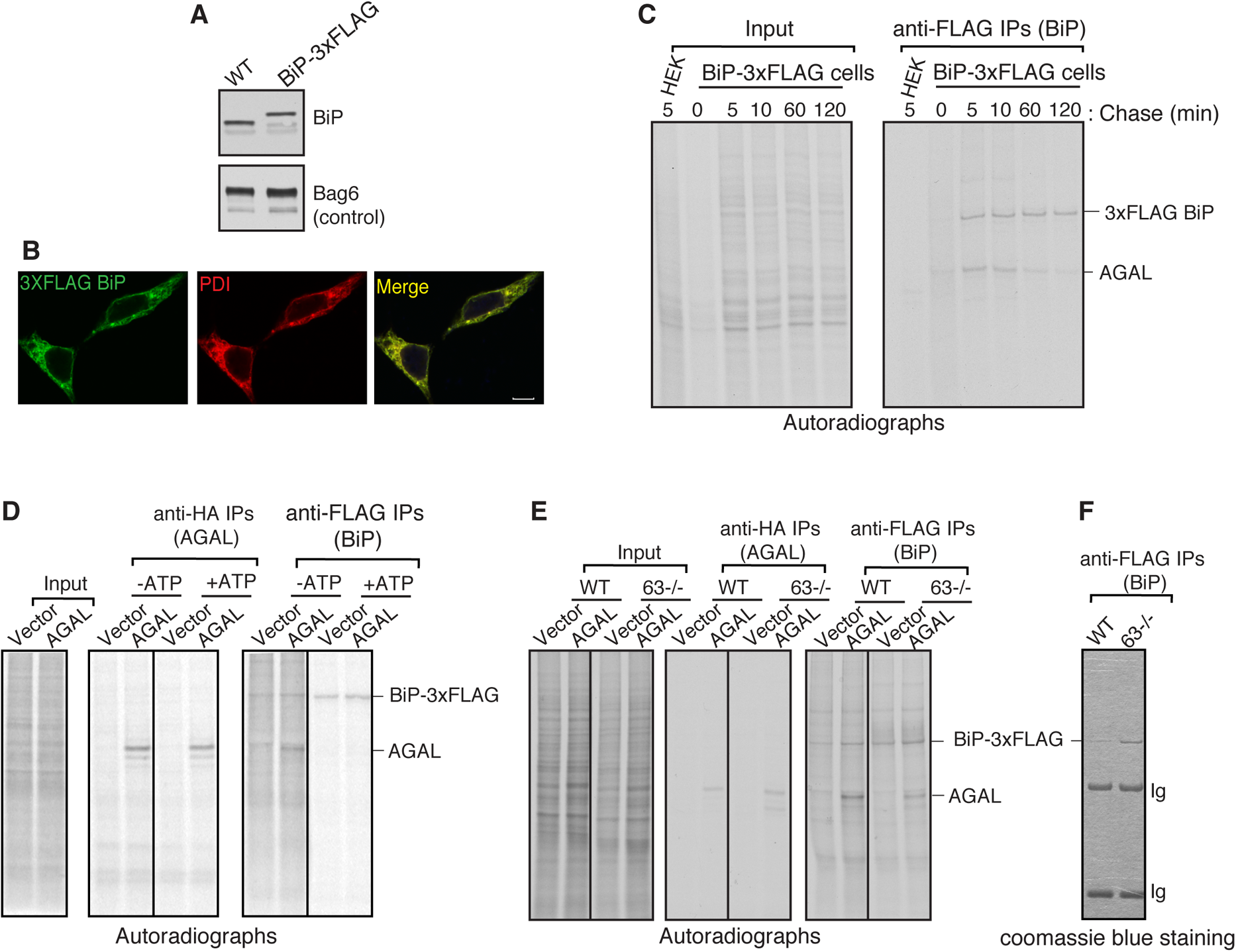
Sec63 mediates BiP binding to translocating nascent chains in the ER. (A) WT HEK293 cells or HEK293 cells that contain a 3xFLAG tag at the C-terminus of endogenous BiP were analyzed by immunoblotting with an anti-BiP antibody. Bag6 serves as a loading control. (B) WT HEK293 and BiP-3xFLAG cell lines were immunostained with anti-FLAG for BiP and anti-PDI antibodies and imaged using a confocal microscope. Scale bars are 10 μm. (C) Cells expressing AGAL-HA were radiolabeled for 2 minutes and chased for the indicated time points. The cell lysates were immunoprecipitated with anti-FLAG antibodies for BiP and analyzed by autoradiography. (D) BiP-3xFLAG HEK293 or BiP-3xFLAG HEK293 Sec63−/− cells were transfected with either empty vector or AGAL-HA. Cells were radiolabeled for 2 minutes, lysed with or without ATP, and immunoprecipitated with anti-FLAG beads before analyzing by autoradiography. (E) BiP-3xFLAG HEK293 or BiP-3xFLAG HEK293 Sec63−/− cells were transfected and radiolabeled as in D. Radiolabeled cells were lysed and immunoprecipitated with either anti-FLAG antibodies for BiP or anti-HA antibodies for AGAL and analyzed by autoradiography. (F) A coomassie stained gel showing immunoprecipitated 3xFLAG tagged endogenous BiP from either BiP-3xFLAG HEK293 cells or BiP-3xFLAG HEK293 Sec63−/− cells. Ig represents immunoglobulins.

To determine if BiP binding to AGAL relies on Sec63, we generated Sec63 knockout in BiP-3xFLAG HEK293 cells using CRISPR/Cas9 (Fig. S6A). Consistent with our model, BiP binding to AGAL was markedly reduced in Sec63−/− cells, although about two-fold more BiP was immunoprecipitated in Sec63−/− cells than WT cells due to BiP upregulation in Sec63−/− cells (Fig. 5E and 5F). Lastly, we tested whether BiP binding to nascent chains of AGAL containing strong SSs also depends on Sec63. To our surprise, similar to AGAL, we found nascent chains of Prl-AGAL and 3L-AGAL also relied on Sec63 activity for binding to BiP, although they were translocated independent of Sec63 (Fig. S6B). This result implies that Sec63 is also required for mediating BiP binding to these nascent chains after their translocation into the ER. Considered together, these results suggest that AGAL nascent chains depend on Sec63 activity for their translocation into the ER as well as their binding to BiP. However, AGAL nascent chains containing strong SSs translocate independent of Sec63 activity but still requires Sec63 for BiP binding after their translocation into the ER lumen.

### BiP-dependent protein translocation is coupled with protein folding in the ER

We hypothesized that BiP binding to AGAL nascent chains via Sec63 may not only facilitate their translocation into the ER but also promote their folding and maturation in the ER. To test this, we monitored the secretion of AGAL since properly folded AGAL is secreted into the extracellular medium upon overexpression (Ioannou et al., 1992). Pulse-chase analysis revealed that intracellular levels of both AGAL and Prl-AGAL in WT cells were reduced during the chase period with a concomitant increase in their extracellular levels (Fig. 6A). By contrast, the secretion of both AGAL and Prl-AGAL was about two-fold reduced in Sec63−/− cells (Fig. 6A). The non-translocated/unglycosylated population of AGAL and Prl-AGAL were degraded by proteasomes in Sec63−/− cells since they were accumulated upon treatment with the proteasomal inhibitor MG132 (Fig. 6A). Moreover, AGAL carrying either a weak or strong SS showed a similar *α*-galactosidase activity in WT cells, but the activity was significantly reduced for all AGAL variants in Sec63−/− cells (Fig. S7A). Reduced activity and secretion in Sec63−/− cells could be attributed to the weak SS of AGAL that relies on Sec63 for efficient translocation into the ER. However, these effects were also observed for even AGAL carrying a strong SS that translocated into the ER mostly independent of Sec63. This indicates that AGAL with a strong SS is likely misfolded after translocation into the ER due to its inefficient binding to BiP in Sec63−/− cells (Fig. S6B).

**Figure 6.**
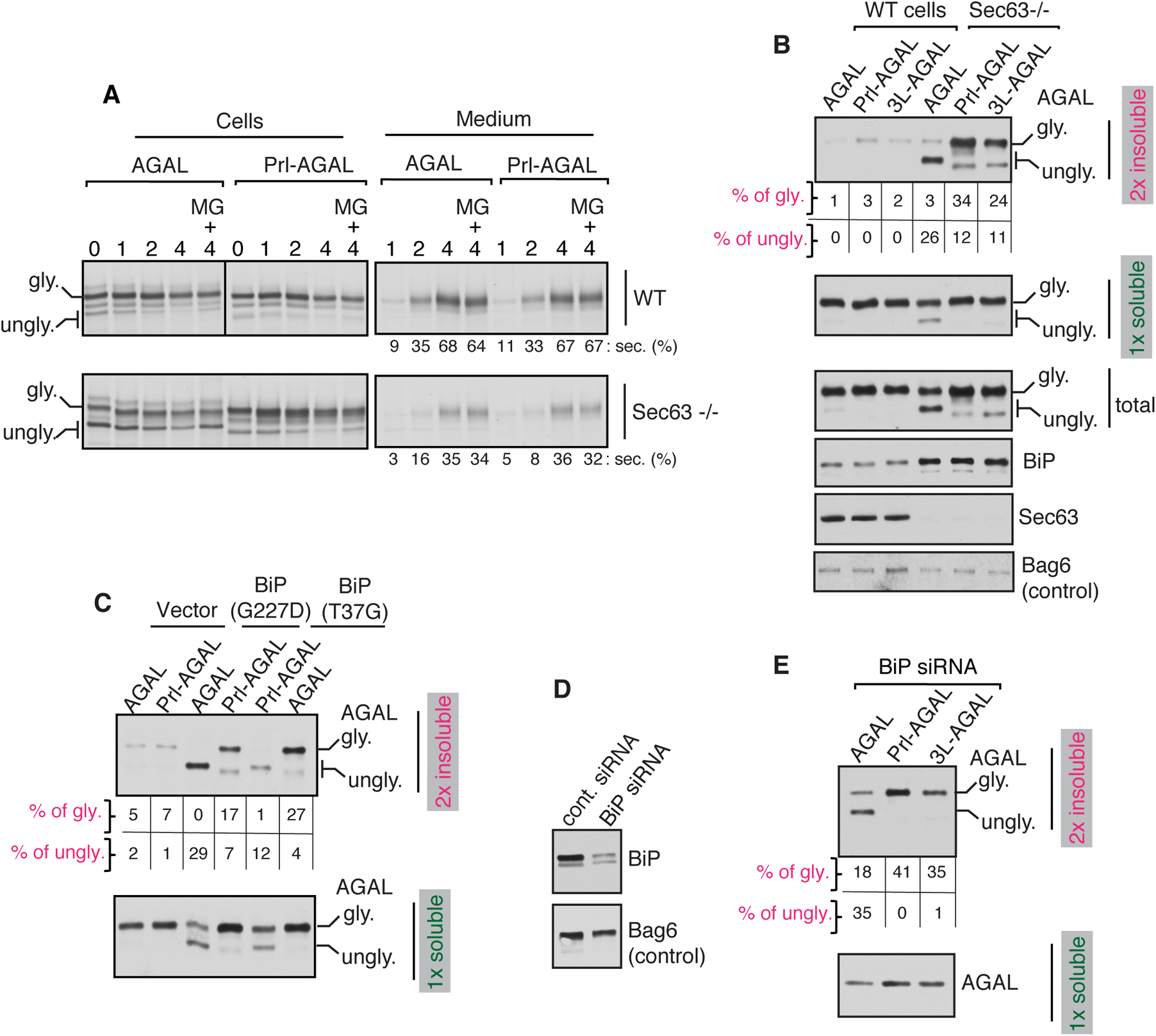
Weak signal sequences sense BiP deficiency and prevents protein misfolding in the ER. (A) WT or Sec63−/− cells expressing AGAL or Prl-AGAL were radiolabeled for 30min and chased in the presence or absence of 20μM MG132 for the indicated time points. Both cells and medium were collected at each time point and analyzed by autoradiography after immunoprecipitation with an anti-HA antibody for AGAL. Secreted AGAL is given as percentage of total signals from both secreted and intracellular bands. “gly” indicates glycosylated forms. “ungly” denotes unglycosylated forms with uncleaved SSs. (B) Cells expressing the indicated constructs were lysed in NP40 buffer, and the aliquots of total lysates and detergent soluble fractions were analyzed by immunoblotting. The detergent-insoluble fractions were lysed in SDS sample buffer and analyzed by immunoblotting. BiP, Sec63, and Bag6 were blotted from soluble fractions. The percent translocated/glycosylated protein values were calculated by insoluble glycosylated signal divided by the sum of glycosylated and unglycosylated signals from both soluble and insoluble fractions. The percent of non-translocated/unglycosylated form was calculated by insoluble unglycosylated signal divided by the sum of glycosylated and unglycosylated signals from both soluble and insoluble fractions. (C) HEK293 cells were co-transfected with the indicated AGAL constructs and vector, BiP (G227D), or BiP (T37G). Cells were treated with 10μM MG132 for 6h before harvesting and analyzing as in B. (D) HEK293 cells were treated with either control siRNA or BiP siRNA and analyzed by immunoblotting for the indicated antigens. (E) BiP siRNA treated HEK293 cells were transfected with the AGAL constructs and harvested after treating the cells with 10μM MG132 for 6h. Cells were lysed and analyzed as in B.

To determine if AGAL carrying a strong SS is misfolded after translocation into the ER of Sec63−/− cells, we separated the detergent soluble fraction from the insoluble fraction, an indicator of misfolded/aggregated proteins. We found a subtle difference between AGAL variants in WT cells. By contrast, a significant fraction of translocated/glycosylated form of Prl-AGAL and 3L-AGAL was sedimented as detergent-insoluble fractions in Sec63−/− cells, supporting our notion that AGAL carrying a strong SS are misfolded and aggregated after translocation into the ER lumen (Fig. 6B). To our surprise, most of the translocated/glycosylated form of AGAL moved to the soluble fraction, but non-translocated/unglycosylated AGAL with the uncleaved SS sedimented as detergent-insoluble aggregates (Fig. 6B and S7B). This result implies that the AGAL population that was translocated even in the absence of Sec63 could be properly folded and matured in Sec63−/− cells. We reasoned that this is due to the upregulation of BiP in Sec63−/− cells, which could partly promote translocation as well as folding of AGAL nascent chains. Sec63-independent but BiP dependent translocation of AGAL in Sec63−/− was supported by the observation that overexpression of BiP ATPase mutants significantly impaired translocation of AGAL in Sec63−/− cells with little effects on the translocation of AGAL with a strong SS (Fig. S7C). Thus, our results suggest that strictly BiP-dependent translocation of AGAL in Sec63−/− ensures subsequent folding and maturation of AGAL in the ER, whereas BiP-independent translocation of AGAL carrying a strong SS is at high risk for misfolding after translocation into the ER of Sec63−/− cells.

Our data thus far suggest a model in which proteins with weak SSs can sense BiP availability in the ER lumen and translocate only upon binding to BiP molecules, which also subsequently assist with protein folding and maturation. In the absence of BiP binding, however, these proteins are synthesized in the cytosol and degraded by proteasomes. To test this model, we co-expressed AGAL variants with BiP mutants (G227D and T37G) that can bind substrates but fail to release them upon ATP binding (Wei et al., 1995). AGAL translocation was markedly reduced in cells expressing BiP mutants and that primarily the non-translocated/unglycosylated AGAL was found in the insoluble fraction (Fig. 6C). However, the population of AGAL that translocated into the ER was completely soluble as shown by the presence of glycosylated form only in the soluble fraction. This result suggests that AGAL either translocates and folds well by binding to active BiP molecules or it does not translocate into the ER when it failed to bind active BiP molecules. Conversely, Prl-AGAL was efficiently translocated/glycosylated irrespective of the BiP functional status in the ER, and thus a significant fraction of Prl-AGAL is misfolded after translocation into the ER and moved to the insoluble fraction (Fig. 6C). Furthermore, our in vitro experiment demonstrated that the translocation of AGAL nascent chain was significantly inhibited in microsomes depleted of luminal proteins compared to that of AGAL containing a strong SS (Fig. S7D).

We further examined whether the weak SS of AGAL can sense BiP availability in the ER by depleting BiP in cells using siRNA (Fig. 6D). The translocation of AGAL but not Prl-AGAL and 3L-AGAL was impaired in BiP depleted cells, which resembles the result seen in Sec63−/− cells (Fig. S7B and S7E). In BiP depleted cells, primarily the non-translocated/unglycosylated form of AGAL moved to the insoluble fraction, whereas a significant population of translocated/glycosylated form of Prl-AGAL and 3L-AGAL went to insoluble fractions (Fig. 6E). These results suggest that the weak SS of AGAL can respond to BiP deficiency by not entering the ER where they would misfold. We then investigated whether the weak SS plays a role in preventing misfolding of AGAL under ER stress conditions, which is known to reduce BiP availability due to the increased load of misfolded proteins in the ER (Kang et al., 2006; Vitale et al., 2019). Indeed, a fraction of Prl-AGAL and 3L-AGAL but not AGAL were readily misfolded and aggregated during ER stress-induced with DTT (Fig. S7F). Collectively, our data suggest that proteins with weak SSs can sense BiP chaperone deficiency in the ER and respond by aborting translocation into the ER to prevent misfolding and aggregation inside the ER.

## Discussion

In addition to targeting roles (Blobel and Dobberstein, 1975), our study presented here suggest that SSs also contain information for protein folding in the ER by coupling protein translocation with the availability of chaperone binding. Our data show that marginally hydrophobic (or weak) SSs containing proteins cause a translocation pause at the Sec61 translocon to recruit luminal BiP chaperone via Sec63 for efficient folding in the ER (Figure 7). This translocation pausing mechanism also allows aborting translocation under conditions of BiP deficiency to prevent protein misfolding and aggregation in the ER. Thus, our studies suggest that evolution has exploited diversity in SSs to recruit selective luminal chaperones for protein folding and maturation of secretory and membrane proteins.

**Figure 7.**
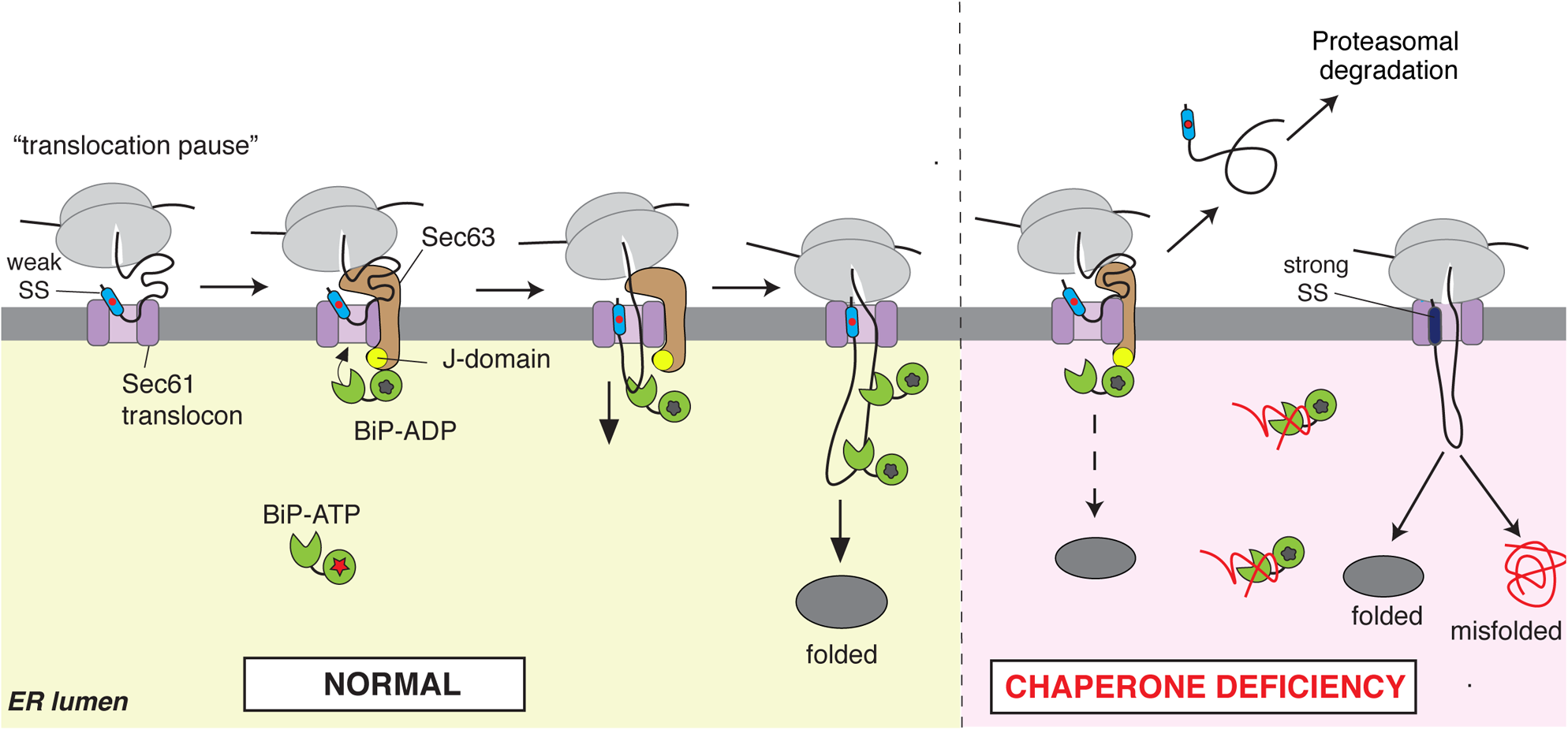
Model for signal sequence-dependent protein folding in the ER. The ribosome-associated nascent chain containing a weak signal sequence (SS) is efficiently recruited to the ER membrane but loosely engages the Sec61 translocon. This causes a pause in the initial translocation of ~160 amino acids in length that accumulates on the cytosolic side of the membrane. Sec63 is likely recruited to the site by sensing the accumulated nascent chains on the cytosolic side of the Sec61 translocon and activates ATPase activity of BiP through its luminal J-domain. Now, ADP-bound BiP can bind onto nascent chains and facilitate their translocation into the ER lumen, thus ensuring subsequent BiP-assisted folding of nascent chains in the ER. Under conditions of BiP deficiency, nascent chains with weak SSs still depend on BiP for translocation, and thus their BiP-assisted folding is guaranteed. However, if nascent chains fail to engage luminal BiP, their translocation is aborted, and they are mislocalized in the cytosol and eliminated by proteasomal degradation. Conversely, even under conditions of BiP deficiency, nascent chains with strong SSs are efficiently translocated in a BiP-independent manner and thus are at a high risk of misfolding and aggregating after translocation into the ER.

Our experiments with model WRB substrates demonstrate that marginally hydrophobic C-terminal TMDs carrying longer C-terminal tails are transiently retained at the Sec61 translocon and are eventually released into the ER lumen. Our data indicate that Sec63 monitors the Sec61 translocon status and directs the clearance of nascent chains transiently retained at the translocon. The key function of Sec63 is to activate the luminal chaperone BiP ATPase through its J-domain so that ADP-bound BiP can bind and release the translocon-retained nascent chain into the ER lumen (Fig. 7). Our findings could explain why the translocation of the C-terminus of membrane proteins is impaired upon disrupting the Sec62/63 complex in yeast (Jung et al., 2014). Our “substrate-trapping” proteomic approach identified numerous secretory and membrane proteins that are transiently retained at the Sec61 translocon complex. Although our data analyzing AGAL and CD44 reveal that marginally hydrophobic SSs plays an important role in determining Sec63 dependency, other SS features such as length and flanking amino acids might also contribute to transient retention of nascent chains at the Sec61 translocon and thus recruiting Sec63/BiP (Schorr et al., 2020b; Ziska et al., 2019).

The role of Sec63 in the post-translational translocation pathway is well established, however, its function in the co-translational protein translocation remains unclear due to the fact that Sec63 and ribosome share the same binding site in the Sec61 translocon (Brodsky et al., 1995; Conti et al., 2015; Itskanov and Park, 2019; Jung et al., 2014; Lang et al., 2012; Matlack et al., 1999; Misselwitz et al., 1998; Wu et al., 2019; Young et al., 2001). Three of our observations suggest that Sec63’s substrates, which we identified here, use the co-translational protein translocation pathway. First, our Sec63 clients are larger than 160 amino acids in length, which appear to be a limitation for post-translational protein translocation in mammalian cells (Lakkaraju et al., 2012). Second, our *in vitro* experiment shows that the Sec63 substrate AGAL could not be post-translationally translocated into the ER. Third, AGAL was efficiently translocated and glycosylated even when it is still attached to the ribosome. Our data also raises the intriguing question of how Sec63 and ribosome can bind to the Sec61 translocon at the same time as they share the same binding site. Our protease protection data suggest that the translocation of nascent chain containing a weak SS is initially paused ~160 amino acids in length and is accumulated on the cytosolic side of the membrane (Fig. 7). This is presumably by caused by a loose interaction between the ribosome nascent chain carrying a weak SS and the Sec61 translocon. We speculate that Sec63 is recruited to the Sec61 translocon by sensing and binding to the cytosolically accumulated nascent chain (Fig. 7). Once recruited, Sec63 J-domain triggers ATP hydrolysis of BiP ATPase to bind recalcitrant nascent chains entering through the Sec61 translocon, thus facilitating their translocation into the ER. Our findings suggest that cells can compensate the loss of Sec63-mediated recruitment of BiP to translocating nascent chains by upregulating BiP levels in Sec63−/− cells. We predict that once the cytosolically accumulated nascent chain at the Sec61 translocon is pulled into the ER lumen, the translating ribosome may establish a tight complex with the Sec61 translocon by displacing Sec63. Our findings may explain why Sec62/Sec63 is selectively enriched with the Sec61 translocon hosting nascent chains of prion protein, which also has a weak signal sequence (Conti et al., 2015; Kang et al., 2006). Our data suggest that Sec62 has a less significant role in the translocation of AGAL, however, additional substrates need to be analyzed to determine its role in translocation and folding of secretory and membrane proteins.

Despite inefficiency in mediating protein translocation, weak SSs are commonly found and conserved across all mammals. One important function for weak SSs is that they can selectively impair the translocation of their mature domains into the ER during ER stress conditions (Kang et al., 2006). Earlier studies have also shown that the nature of the interaction between a SS and the Sec61 translocon can influence downstream events such as the timing of SS cleavage and N-linked glycosylation (Rutkowski et al., 2001; Rutkowski et al., 2003), which in turn may direct protein maturation (Lingappa et al., 2002). Our study now suggests that Sec63/BiP-mediated release of initial translocation pausing of nascent chains carrying weak SSs is coupled with the BiP-assisted protein folding and maturation in the ER. This mechanism may be particularly important for the folding of AGAL since it contains the hydrophobic N-terminal region right after the signal sequence that is buried into the structure, which is also the hotspot for genetic mutations causing Fabry disease (Garman, 2007). When functional BiP is scarce in the ER under conditions such as stress and starvation, the translocation of substrates like AGAL is prevented and thus misfolding and aggregation in the ER lumen is avoided. Although the detergent soluble assay used here provides evidence of protein folding and misfolding, it does not provide quantitative information. New experimental approaches such as a folding sensitive fluorescence reporter are required to quantitatively determine signal sequence-dependent folding in live cells. The non-translocated substrates are likely eliminated by the Bag6-mediated cytosolic quality control pathway (Hessa et al., 2011).

Our data show that increasing the hydrophobicity of weak SSs is sufficient to bypass the BiP-dependent translocation, but translocated nascent chains are at risk of misfolding and aggregation under conditions of BiP deficiency. Our findings may explain why about 40% of the analyzed yeast secretome use the Sec63/Sec62-mediated post-translational translocation pathway (Ast et al., 2013), which likely reflects their dependency on BiP for folding and maturation. We predict that the recruitment of BiP may occur at any time during translocation as long as nascent chains contain a translocon retention element such as marginally hydrophobic sequences. Consistent with this notion, earlier studies have observed translocation pausing events in which the nascent chain transiently exposes to the cytosol during the translocation of apolipoprotein B and immunoglobulin G (Hegde and Lingappa, 1996; Rutkowski et al., 2001). We suggest that these translocation pausing events may promote maturation of proteins by recruiting specific luminal chaperones to nascent chains.

We propose that the translocon-associated accessory proteins may function as a hub to recruit specific luminal chaperones and enzymes to bind onto translocating nascent chains. In addition to marginally hydrophobic SSs that recruit BiP through Sec63, we propose other features in SSs may also modulate translocon accessory proteins and their downstream chaperones. This notion is supported by the recent findings that glycine and proline-rich SSs containing proteins depend on the translocon-associated protein (TRAP) complex for their efficient translocation into the ER (Nguyen et al., 2018). Curiously, the endogenous TRAP complex was recently shown to form a complex with Calnexin chaperone (Lee et al., 2020), suggesting a potential role in chaperone selection and glycoprotein folding. It is tempting suggest that multiple cellular translocation pathways (Aviram and Schuldiner, 2017) in cells may have evolved to meet the challenges of folding the complex secretome by recruiting protein-specific chaperones.

Earlier studies have shown that mutations in the Sec63 gene can cause autosomal dominant polycystic kidney diseases (PKD) due to defects in protein maturation and cell surface localization of polycystin 1 and presumably other proteins (Besse et al., 2017; Davila et al., 2004). Around one-third of our Sec63 clients have been previously implicated in human genetic disorders, including many lysosomal storage disorders and PKD. This suggests that these proteins are particularly vulnerable to misfolding and aggregation due to genetic mutations. In light of our findings, selective induction of BiP by small molecules, such as AA147 (Plate et al., 2016), may be a potential therapeutic approach to treat these diseases. Furthermore, our studies may guide the design of customized SSs to produce a growing number of therapeutic recombinant secretory proteins, including enzymes, growth factors, and antibodies.

## Materials and Methods

### DNA constructs

Most of the cDNAs used in this study were cloned into pcDNA/FRT/TO vector (Invitrogen). pcDNA/FRT/TO containing WRB-HA-Venus substrates were previously described (Sun and Mariappan, 2020). Mouse Sec63-FLAG (template mouse Sec63 plasmid was a kind gift from Dr. Stefan Somlo, Yale School of Medicine). Sec63 HPD/AAA (H132A/P1133A/D134A) and Sec63 *Δ*230-330 were previously described (Li et al., 2020). Prl-AGAL-FLAG and CD44-FLAG constructs were generated by replacing AGAL SS (1-31aa) and CD44 SS (1-20aa) with bovine prolactin SS (1-31aa) using phosphorylated primers and Phusion Site-Directed Mutagenesis procedure. Since the flanking threonine (T) of prolactin SS that remained after the SS cleavage inhibited AGAL activity, Prl-AGAL-HA used in Fig. 5 and Fig. 6 was created by replacing AGAL SS (1-31aa) with bovine prolactin SS (1-30 aa) using the above procedure. 3L-AGAL and 3L-CD44 constructs were created using Pfu polymerase (Agilent Technologies) based site-directed mutagenesis method. AGAL-prolactin-FLAG was cloned by replacing bovine prolactin SS (1-30) with AGAL SS (1-31) using the above procedure. BiP mutants (G227D and T37G) were introduced into pcDNA Rat BiP using the above site-directed mutagenesis method. The coding sequences of all constructs were verified by sequencing (Yale Keck DNA Sequencing).

### Antibodies and reagents

Antibodies used for immunoblotting are as follows: Rat α-FLAG L5 (BioLegend #12775), Rabbit α-HA, Rabbit α-GFP, Rabbit α-Sec63, Rabbit α-Sec63, Rabbit α-Sec61β, Rabbit α-BAG6, and Rabbit α-Sec61α are a gift from Dr. Ramanujan Hegde. Rabbit α-BiP (Proteintech #11587-1-AP), Mouse α-HA (Biolegend #901513), Mouse α-PDI (Affinity Bioreagents #MA3-018), Goat α-Rat-HRP (Cell Signaling, #7077), Goat α-Mouse-HRP (Jackson ImmunoResearch, #115-035-003), Goat α-Rabbit-HRP (Jackson ImmunoResearch, #111-035-003), Goat anti-rat HRP (Cell signaling, #7077S) Goat α-RAT IgG-Cy2 (Jackson ImmunoResearch # 112-225-167), Goat α-Mouse IgG-Alexa657 (Invitrogen # A-21235). Beads were purchased as follows: Strep-Tactin XT beads (IBA, Germany #2-4010-010), Rat anti-FLAG L5 affinity gel (Biolegend, #651503), Protein A agarose (Repligen, #CA-PRI-0100), Mouse anti-HA magnetic beads (Pierce, #88837), Poly L-lysine (Peptides International, # OKK-3056). Rabbit Reticulocyte Lysate was purchased from Green Hectares (Ph:1-800-GHLYSAT). Detergents were purchased as follows: Digitonin (EMD Millipore, Billerica, MA), Triton X-100 (Thermo Fisher), Sodium Deoxycholate (Sigma), and Sodium Dodecyl Sulfate (Sigma), Saponin (Sigma), 37% Formaldehyde (Avantor), and Hoechst 33342 stain (Cell Signaling # 4082S), Tween 20 (American Bioanalytical).

### Cell culture and CRISPR/Cas9 genome editing

HEK293-Flp-In T-Rex cells (Invitrogen) were cultured in high glucose DMEM containing 10% FBS at 5% CO_2_. The knockout cell lines of Sec63 or Sec62 as well as 2xStrep-Sec61α,cells were previously described (Li et al., 2020). To engineer chromosomal BiP 3xFLAG HEK293 cells, the BiP gRNA sequence (5′ CCTCTTCACCAGTTGGGGG 3′) was cloned into the gRNA expression vector (Mali et al., 2013). The single-strand DNA oligonucleotide homology-directed repair (HDR) sequence (CTGGAAGAAATTGTTCAACCAATTATCAGCAAACTCTATGGAAGTGCAGGCCCTCCCCCAgccGACTACAAGGACCACGACGGCGATTATAAGGATCACGACATCGACTACAAAGACGACGATGACAAGgcaACTGGTGAAGAGGATACAGCAGAAAAAGATGAGTTGTAGACACTGATCTGCTAGTGCTGTAA) was synthesized (IDT) for inserting 3xFLAG-tag into the C terminus of BiP encoding gene in HEK293 cells. While the sequence indicated in blue encodes for 3xFLAG, green denotes the C-terminus KDEL peptide encoding sequence. Pink indicates the stop codon. 300 pmol of HDR oligonucleotide was electroporated into one million HEK293 Flp-In T-Rex cells along with 2.5 μg each of pSpCas9(BB)-2A-Puro and BiP gRNA plasmid (Amaxa kit R, program A-24; Lonza). Immediately after electroporation, cells were plated in a 6 well plate. After 24 h of electroporation, the expression of Cas9 was selected by puromycin treatment (2.5 μg/ml) for 48 h. Most cells were died of electroporation and puromycin treatment, but about 5% survived cells were grown for a week without puromycin. Cells were replated at 0.5 cell/well in 96 well plates and expanded for 3 weeks. Individual clones were examined for the presence of 3xFLAG BiP by immunoblotting with anti-BiP antibodies. Positive clones containing 3xFLAG BiP exhibited a slower migrating band compared to BiP present in control HEK293 cells.

### Immunoprecipitations

To test the interaction between WRB-Venus substrates and the Sec61 translocon complex, the chromosomally Sec61α-2x Strep-tagged HEK293 cells were plated on a poly-lysine (0.15mg/ml) coated 6 well plates at 0.6 million/well. The cells were transiently transfected with 2μg WRB-Venus substrates using 5μL of Lipofectamine 2000 and treated with 200 ng/ml doxycycline to induce protein expression. After 24 hours of transfection, cells were harvested in 1XPBS and centrifuged for 2 min at 10,000g. The cell pellet was lysed in 200μl of lysis buffer (50 mM Tris pH 8.0, 150 mM NaCl, 5 mM MgAc, 2% digitonin, and 1X Roche EDTA-free protease inhibitor cocktail) by incubating on ice for 30 min and then diluted to 1ml with the lysis buffer containing 0.1% digitonin. The 5% digitonin (Millipore) stock was boiled for 5 min just before adding into the lysis buffer to avoid digitonin precipitation during IPs. The supernatants were collected by centrifugation at 15,000g for 15 min at 4°C. For co-immunoprecipitation, the supernatant was rotated with StrepTactin XT beads (IBA, Germany) for 2h at 4°C. The beads were washed 3 times with 1 ml of wash buffer (50 mM Tris pH 8.0, 150 mM NaCl, 5 mM MgAc, 0.1% digitonin). The bound material was eluted from the beads by directly boiling in 50 μl of 2x SDS sample buffer for 5 min and analyzed by immunoblotting.

To test the interaction between Sec63-FLAG and Sec61 translocon shown in Fig. S2A, the above digitonin IP procedure was followed but using HEK293 Sec63−/− cells stably expressing variants of Sec63-FLAG. 20ng/mL doxycycline was used to induce the protein expression for 24 h before harvesting for IPs.

### Metabolic labeling and immunoprecipitation

Indicated cells (0.6 million cells/well) were plated on poly-lysine coated (0.15mg/mL) 6-well plates and transiently transfected with 2μg indicated plasmids. The expression of indicated proteins was induced with doxycycline (200ng/mL) for 24 hours. The cells were starved with Met/Cys-free medium including 10% dialyzed FBS and doxycycline for 30 min. Cells were then labeled with 80μCi/mL Express-^35^S protein labeling Mix (NEG772007MC) for 30 min. The labeled cells were directly harvested in 1ml of RIPA buffer including 1X EDTA-free protease inhibitor cocktail (Roche). The lysate was cleared by centrifugation at 20,000g for 15 min. The supernatant was mixed with the unconjugated or antibodies conjugated to beads indicated in the figure legends for 1.5 h in the cold room. In the case of unconjugated anti-rabbit antibodies, 20ul slurry of protein-A agarose was added to samples and incubated for another 1 hour. The beads were washed three times with 1 ml of RIPA buffer, and proteins were eluted with 50ul of 2X SDS sample buffer and run on 7.5% or 10% Tris-Tricine SDS-PAGE gels. The gels were dried and analyzed by autoradiography.

For isolating endogenous substrates transiently retained at the Sec61 tranlocon, Sec63−/− and Sec63−/− cells complemented with WT Sec63, Sec63 J-mutant (HPD/AAA) or the translocon interaction defective Sec63 mutant (Δ230-300) were labeled as described above. The labeled cells were washed and harvested 1XPBS and centrifuged at 10,000g for 2 min. The cell pellets were lysed in 200μL of lysis buffer (50 mM Tris pH 8.0, 150 mM NaCl, 5 mM MgAc, 2% digitonin, and 1X Roche protease inhibitor cocktail) by incubating on ice for 30 min and then diluted to 1ml with the lysis buffer containing 0.1% digitonin. The supernatant was collected by centrifugation at 15,000g for 15 min and incubated with Rat anti-FLAG beads (BioLegend) for 2h at 4°C. The beads were washed 3 times with 1 ml of wash buffer (50 mM Tris pH 8.0, 150 mM NaCl, 5 mM MgAc, 0.1% digitonin). The bound material was eluted from beads by directly boiling in 50 μl of 2x SDS sample buffer for 5 min and analyzed by SDS-PAGE and autoradiography.

For pulse-chase analysis, transfected cells were labeled as above for 30 min and chased in complete DMEM medium supplemented with label-free 2mM Methionine and 2mM Cysteine for indicated time points. The cells were directly harvested in RIPA buffer including 1X EDTA free protease inhibitor cocktail (Roche) and cleared by centrifugation at 15,000g for 15 min. The supernatant was incubated with the indicated antibody conjugated to beads for 2 h. For checking the secretion of AGAL or Prl-AGAL, the medium was collected at each time point, adjusted to 50 mM Tris pH 8.0, 150 mM NaCl, 1% Triton X-100, and immunoprecipitated and analyzed as above.

### Endogenous BiP-3xFLAG immunoprecipitation

BiP-3xFLAG HEK293 cells (0.9 million cells/well) were plated on poly-lysine coated (6-well plates and transiently transfected with 2μg of indicated plasmids and induced with doxycycline (200ng/mL) for 24 hours. The cells were starved with Met/Cys-free medium including 2.5% dialyzed FBS and doxycycline for 45 min. Cells were then labeled with 90 μCi/mL Express-^35^S protein labeling Mix (NEG772007MC) for 2 min at 37°C. The labeled cells were immediately placed on ice and harvested in 1ml of ice-cold 1X PBS and centrifuged at 10,000g for 2 min. Cell pellets were solubilized in 200 ul of lysis buffer (50 mM Tris, pH 7.5, 160 mM NaCl, 1% Digitonin, 1mM CaCl2, Apyrase (10 U/mL)) for 30 min on ice. The lysate was diluted to 1ml with lysis buffer containing 0.1% Digitonin before centrifugation at 20,000g for 15 min. While 800ul of supernatant was incubated with 12 ul of mouse anti-FLAG beads (Sigma), the remaining 200ul was diluted to 800ul with NP40 buffer and incubated 15 ul mouse anti-HA magnetic beads (Thermo Scientific). After 90 min of incubation, anti-FLAG beads were washed 3x with 1ml of lysis buffer containing 0.1% Digitonin but anti-HA magnetic beads were washed with 3x with 1ml of NP40 buffer. The washed beads were boiled with 50ul of 2x SDS samples buffer and analyzed by 7.5% Tris-Tricine SDS PAGE and autoradiography. For the 2^nd^ IP, anti-FLAG immunoprecipitants (~40 ul in 2x SDS sample buffer) were diluted to 1ml with Triton buffer (50 mM Tris, pH 8, 150 mM NaCl, 1% Triton X-100) and incubated with 15ul of anti-HA magnetic beads for 90 min. The beads were washed 3x with 1ml of Triton buffer and analyzed as above.

### Affinity purification of translocon-retained endogenous substrates and mass spectrometry

Three different HEK293 cell lines (Sec63−/−, Sec63−/− stably complemented with WT Sec63-FLAG, Sec63−/− complemented with Sec63-FLAG J-mutant (HPD/AAA) were plated in 10cm plates at 10 million cells/plate. Protein expression was induced by 20ng/mL doxycycline. After 24 h of induction, the cells were transferred from one 10cm plate to two 15cm plates for each cell line including 20ng/mL doxycycline. After culturing two days, cells at ~100% confluence were treated with 100ug/mL cycloheximide for 5 min. The cells were then collected using ice-cold 1XPBS and centrifuged at 1500 g for 10 min to obtain the cell pellets. The cell pellets were resuspended in lysis buffer (50mM HEPES pH 7.4, 100mM KAc, 50mM NaCl, 10mM MgAc, 1% digitonin, 1X Roche protease inhibitor cocktail, 1mM DTT, 100ug/mL cycloheximide). The lysates were further homogenized using a dounce by passing through manually for 10 times. The lysates were then centrifuged at 35,000g for 30 min at 4°C in an MLA80 rotor. The supernatant was incubated with 300μL of pre-washed mouse anti-FLAG beads (Sigma) slurry for 2 hours at 4°C. The beads were washed for 6 times with 5 mL of wash buffer (50mM HEPES pH 7.4, 100mM KAC, 50mM NaCl, 10mM MgAC, 0.1% digitonin, 1mM DTT). The samples were eluted with the elution buffer (0.1M Glycine pH2.3 and 0.1% Triton X-100). The eluted materials were TCA precipitated and subjected to mass spectrometry analysis.

The immunoprecipitants were digested with trypsin and analyzed by LC-MS/MS using an LTQ Orbitrap Elite equipped with a Waters nanoACQUITY ultra-performance liquid chromatography (UPLC) system at the Mass Spectrometry & Proteomics Resource of the W.M. Keck Foundation Biotechnology Resource Laboratory, Yale University. Proteins were searched against the homo sapiens SwissProt protein database using the Mascot Search Engine (Matrix Science, LLC, Boston, MA; version. 2.6.0). Scaffold program (version Scaffold_4.8.7; Proteome Software, Portland, OR) was used to validate MS/MS-based peptide and protein identifications. Scaffold program was also used to generate normalized spectral counts for each sample. In the supplemental Table 1, we listed protein hits from all three samples that have a protein threshold of 5.0% FDR, at least 2 peptides, and a peptide threshold of 95%. In the supplemental Table 2, we listed protein hits of secretory/membrane proteins from Sec63 J-mutant that are at least 2-fold enriched relative to WT Sec63 and 5-fold enriched relative to negative control Sec63−/− cells. The information of SSs or the 1^st^ TMD sequences, subcellular localization, and association with human diseases were extracted from UniProt.Org. In the supplemental Table 3, we listed secretory/membrane protein hits from WT Sec63 that are at least 2-fold enriched relative to Sec63 J-mutant and 5-fold enriched relative to negative control Sec63−/− cells. The hydrophobic region of a signal sequence shown in Fig. 3D, E was defined using the MacVector program by selecting the Kyte-Doolittle hydrophilicity option in windows of 3-19 amino acids. The defined hydrophobic region was analyzed by grand average hydropathy (GRAVY) to obtain a hydrophobicity score.

### Endo H treatment

Indicated cell lines (0.15 million cells/well) were plated on poly-lysine (0.15mg/mL) coated 24-well plates and transiently transfected with 400ng indicated plasmids. Protein expression was induced with 200ng/mL doxycycline. After 24 h of transfection, the cells were collected in 150μL of 150ul 2x SDS sampled buffer and boiled for 10min. 20ul sample was mixed with 65ul of water, 5ul of 20% Triton X-100, 10 ul of 10X G5 buffer (NEB), and 1ul of Endo H HF (NEB). The reactions were incubated for 4h at and were terminated by adding 50μL 5x SDS sample buffer and boiled for 5 min before analyzing by immunoblotting. The above procedure was used for Endo H digestion of immunoprecipitated samples that were already boiled in a 2x SDS sample.

### *In vitro* transcription and translation

The open reading frames were PCR amplified from pcDNA/FRT/TO based constructs using a forward primer bearing Sp6 promoter sequence and annealing sequence to the CMV promoter and a reverse primer that anneals to the poly-A region of the vector. For generating ribosome-associated nascent chains (RNCs), PCR products were amplified using the above forward primer, but the reverse primer anneals to 3’ of the template cDNA without the stop codon. In vitro transcription reactions including PCR product, SP6 polymerase (NEB), and RNasin (Promega) were performed at 37°C for 1.5h as described previously (Sharma et al., 2010). The transcripts were translated in the rabbit reticulocyte lysate (RRL) translation system including ^35^S-methionine and with or without rough microsomes. Rough microsomes were either prepared from the canine pancreas as described previously (Walter and Blobel, 1983) or cultured HEK293 or Sec63−/− cells as described previously (Zhang et al., 2013). Transcripts encoding WRB-Venus variants were translated in the presence or absence of HEK or Sec63−/− microsomes for 30 min at 32°C. The translated products (10μL) were denatured with 5x volume of 2x SDS sample buffer and analyzed by SDS-PAGE and autoradiography.

For the co-translational translocation assay, transcripts encoding AGAL-FLAG, Prl-AGAL-FLAG were translated in RRL in the presence or absence of canine pancreas microsomes (CRM) for 30 min at 32°C. The translation products were treated or untreated with 0.5mg/mL Proteinase K (PK) for 60 min on ice. PK digestion was terminated by adding 5 mM PMSF and incubating on ice for 15 min. The resulting samples were transferred into the boiling SDS buffer (100 mM Tris pH 8.0 and 1% SDS) and heated for an additional 5 min. The samples were diluted 10 times with the Triton buffer (50 mM Tris pH 8.0, 150mM NaCl, 1% Triton X-100). The diluted samples were incubated with anti-FLAG beads (Bio-legend) for 2 hours at 4°C. The beads were washed three times with 1ml of Triton buffer and eluted by directly boiling in 50 μl of 2XSDS sample buffer for 5 min and analyzed by SDS-PAGE and autoradiography. For the post-translational translocation assay, transcripts encoding AGAL-FLAG were translated in RRL for 30 min at 32°C. The translation reactions were centrifuged at 70,000 rpm in a TLA120.1 rotor for 30 min at 4°C. The post-ribosomal supernatant was incubated with or without microsomes for 30 min at 32°C. The resulting samples were treated with proteinase K and analyzed as above.

RNCs of indicated lengths were produced from transcripts lacking stop codon in the presence or absence of the indicated microsomes. The resulting translated products were split into two and incubated with or without 0.5mg/mL Proteinase K (PK) for 60 min on ice. The PK digestion was terminated by adding 5 mM PMSF and incubating on ice for 15 min. The samples were transferred into boiling the 2x SDS sample buffer and incubated for an additional 5 min. For RNCs of AGAL or Prl-ss-AGAL, the translated products were analyzed by autoradiography after denaturation and immunoprecipitation with anti-FLAG beads as described above. In some cases, the denatured samples were treated with RNase A for 15 min at 37°C to remove tRNAs from nascent chains before analyzing by SDS PAGE and autoradiography.

Luminal proteins depleted CRM shown in Fig. 6C was prepared as described previously (Kang et al., 2006), 100ul CRM at a concentration of 1eq/ul was diluted with 400ul with membrane buffer (MB: 50 HEPES 7.4, 250mM Sucrose, 2mM MgCl2, 1mM DTT, and 0.075% deoxy Big CHAP (DBC)) for 10min on ice. The detergent permeabilized microsomes were centrifuged for 70,000 rpm for 30min in a TLA100.3 rotor. The resulting pellet was resuspended in 500ul of MB without DBC and centrifuged as above. The final pellet was resuspended in 50ul of MB without DBC and used for in vitro translation reactions as indicated above.

### Membrane recruitment assay

We followed the previously published procedure (Yanagitani et al., 2011) with the following modifications. Transcripts encoding versions of AGAL lacking a stop codon were translated in RRL including ^35^S-methionine in the presence of either trypsin (20ug/ml) digested canine rough microsomes (CRM) (Plumb et al., 2015) or CRM for 20 min at 32°C. 25ul translation reactions were layered on 100ul of 1 M sucrose prepared in 50mM HEPES pH 7.5, 100 mM NaCl, 5mM MgCl2. After sedimentation for 15 min at 20,000×g, 100 ul supernatants were carefully collected and pellets were resuspended in 75ul of 1M sucrose. Aliquots of both supernatants and pellets were analyzed by SDS-PAGE and autoradiography.

### Isolation of detergent-soluble and -insoluble fractions

The indicated cells (0.35 x 10^6^/well) were plated on a poly-lysine coated 12-well plate and transiently transfected with 0.8ug of AGAL-HA or its variants. Protein expression was induced with 200ng/mL doxycycline. After 24 hours of induction, the cells were harvested with 1X PBS and centrifuged to collect the cell pellet. The cell pellet was resuspended with 400μL of NP40 buffer (50 mM Tris, pH 7.5, 150 mM NaCl, 0.5% sodium deoxycholate, and 0.5% NP40) and centrifuged for 10 min at 14,000 rpm at 4°C. The detergent soluble supernatant was collected, and the detergent-insoluble pellet was directly boiled in 200 ul of 2XSDS buffer for 10min. Both soluble and insoluble fractions were analyzed by immunoblotting.

### Immunofluorescence

BiP-3xFLAG HEK293 cells (0.12 × 10^6^) were plated on 12 mm round glass coverslips (Fisher Scientific) coated with 0.15 mg/mL poly-lysine in 24-well plates. The following day, cells were fixed and immunostained as previously described (Sundaram et al., 2017). Rat anti-FLAG antibody (BioLegend) and mouse anti-PDI antibody (Affinity Bioreagents) were used as primary antibodies. Goat anti-mouse Alexa 647 and Goat anti-rat Cy2 (Jackson Immuno Research) were used as secondary antibodies. The cells were imaged on a Leica scanning confocal microscope as previously described (Sundaram et al., 2017).

### α-galactosidase-A activity assay

Cells (0.35 x 10^6^/well) were plated on poly-lysine (0.15mg/mL) coated 12-well plates and transiently transfected with 800ng AGAL-FLAG versions. After 24 hours of transfection, the cells were washed once with 1XPBS and harvested in 1XPBS. After centrifugation at 10000 g for 2 min, the cell pellet was resuspended in the 100μL of NP40 buffer (50 mM Tris, pH 7.5, 150 mM NaCl, 0.5% NP40, and 0.5% deoxycholate) and incubated on ice for 20min. The lysate was centrifuged for 10 min at 14000 g at 4°C. The assay was performed as described previously with the following modifications (Lukas et al., 2016). 50μL of 2mM artificial substrate 4-MU-α-D-galactopyranoside (Sigma) dissolved in 0.06 M phosphate citrate buffer (pH 4.7) was mixed well with 1μL of the supernatant in a black 96 well plate (Greiner, #655076) for fluorescence reading, and incubated for 5min at 37°C. The reaction was terminated by the addition of 200μL 1 M glycine buffer (pH 10.5). The released 4-MU was measured by a fluorescence reader (TECAN) using 360nm excitation filter and 465nm emission filter. The readings were subtracted from empty vector-transfected cells, and AGAL variant activities were normalized to wild type AGAL activity.

### Quantification and statistical analysis

Quantification of bands in immunoblots and autoradiographs were performed using the ImageJ software (NIH). The graphs were created by Prism7, and the error bars reflect the standard error of the mean from three independent experiments.

### Online supplemental material

**Fig. S1** shows the characterization of Sec63-mediated clearance of translocon-retained marginally hydrophobic TMDs. **Fig. S2** demonstrates both J-domain and translocon interacting regions are required for releasing translocon-retained substrates. **Fig. S3** supports the mislocalization of AGAL in the cytosol of Sec63−/− cells. **Fig. S4** shows conserved weak signal sequences determine Sec63 dependency. **Fig. S5** shows the weak signal sequence of AGAL can efficiently target its ribosome-nascent chain to the ER membrane. **Fig. S6** shows Sec63 mediates BiP binding to translocating nascent chains containing both weak and strong signal sequences. **Fig. 7** demonstrates the effect of signal sequences on protein translocation and protein folding. **Table S1** shows proteins identified from Sec63−/−, WT Sec63, or Sec63 J-domain mutant sample. **Table S2** shows secretory and membrane proteins that are enriched in Sec63 J-mutant**. Table S3** shows secretory and transmembrane proteins that are enriched in WT Sec63.

## Acknowledgements

We are grateful to Jean Kanyo and TuKiet Lam from the Yale Keck Protromic facility for assisting with analyzing mass spectrometry data. We thank Jacob Culver and Matthew Jordan for the discussion. We thank Jacob Culver, Sang-Wook Kang, and Zai-Rong Zhang for comments on the manuscript. We are thankful to Fred Gorelick for the comments and edits on the manuscript. This work is funded by NIH grant R01GM117386 (M.M).

The authors declare no competing financial interests.

## Author contributions

SS designed and performed most of the experiments. XL performed imaging and BiP binding and dissociation experiments. MM generated BiP-3xFLAG HEK293 cells, performed BiP interaction studies, and supervised the project. SS and MM wrote the manuscript with inputs from XL.

## Figure Legends

**Table S1. Proteins identified from Sec63−/−, WT Sec63, or Sec63 J-domain mutant sample.** The scaffold program was used to generate normalized spectral counts for each sample. The table shows protein hits from all three samples that have protein threshold of 5.0% FDR, at least 2 peptides, and peptide threshold of 95%.

**Table S2. Secretory and membrane proteins that are enriched in Sec63 J-mutant.** The table shows protein hits from Sec63 J-mutant that is at least 2-fold enriched relative to WT Sec63 and 5-fold enriched relative to negative control Sec63−/− cells. The information of SSs or the 1st TMD sequences, subcellular localization, and association with human diseases were extracted from UniProt.Org. The red color labeled proteins were also found in the list of Sec63 substrate candidates identified by Schorr et al, the FEBS Journal, 2020. Note that the known Sec63 interaction proteins, Sec62 and Sec61β, were manually removed from the list.

**Table S3. Secretory and transmembrane proteins that are enriched in WT Sec63.** The table shows protein hits from Sec63 WT that is at least 2-fold enriched relative to Sec63 J-mutant and 5-fold enriched relative to negative control Sec63−/− cells. The information of subcellular localizations is extracted from UniProt.Org. The red color labeled proteins were also found in the list of Sec63 substrate candidates identified by Schorr et al, the FEBS Journal, 2020. Note that the mitochondrial proteins were manually removed from the list.

## Notes

### Competing Interest Statement

The authors have declared no competing interest.

