## Supplemental Figures for "Signal sequences encode information for protein folding in the endoplasmic reticulum"

### Figure S1

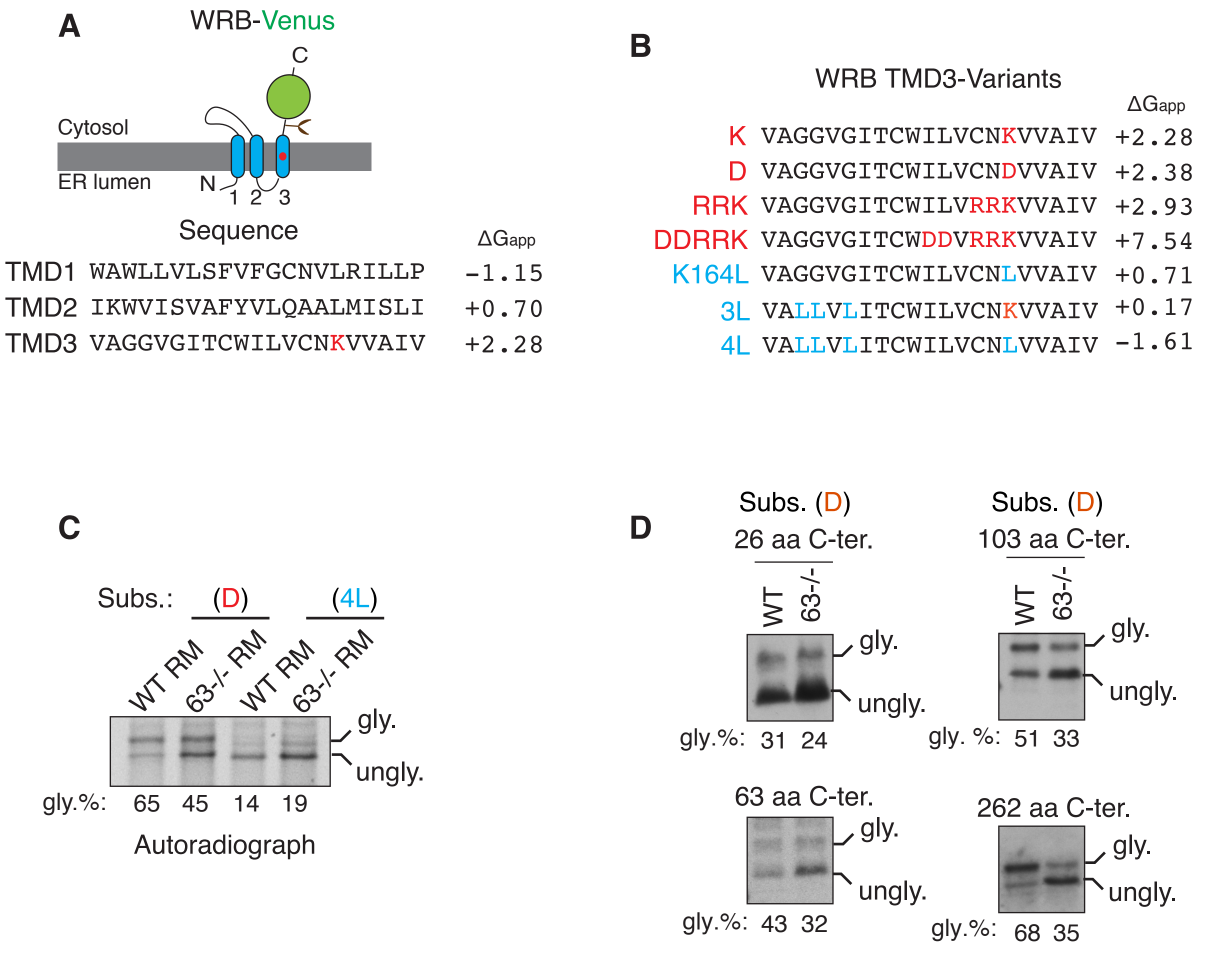


**Figure S1. Characterization of Sec63-mediated clearance of translocon retained marginally hydrophobic TMDs**

(A) A cartoon depicting the predicted topology of WRB fused with the C-terminus Venus tag. The red circle in the C-TMD (3rd TMD) indicates the charged lysine amino acid. The amino acid sequence and free energy prediction for all three TMDs of WRB are indicated. Note that the C-TMD has a high free energy (ΔGapp) value because it contains a positively charged lysine residue.

(B) The C-TMD of WRB-Venus is mutated to either increase the free energy by introducing charged residues as specified in red color or reduce the free energy by introducing hydrophobic leucine residues as indicated in blue color (Sun and Mariappan, 2020). The free energy of all C-TMD variants was predicted as previously described (Hessa et al., 2007).

(C) Transcripts encoding the indicated substrates were translated in rabbit reticulocyte lysate (RRL) in the presence of rough microsomes (RM) prepared from either WT HEK293 or Sec63-/- cells. The reactions were directly analyzed by SDS-PAGE and autoradiography.

(D) WT HEK293 or Sec63-/- cells were transfected with the marginal hydrophobicity substrate (Subs. D) that contains the indicated length of amino acids after their C-TMDs. Cell lysates were immunoprecipitated and analyzed by immunoblotting with anti-HA antibodies.

### Figure S2

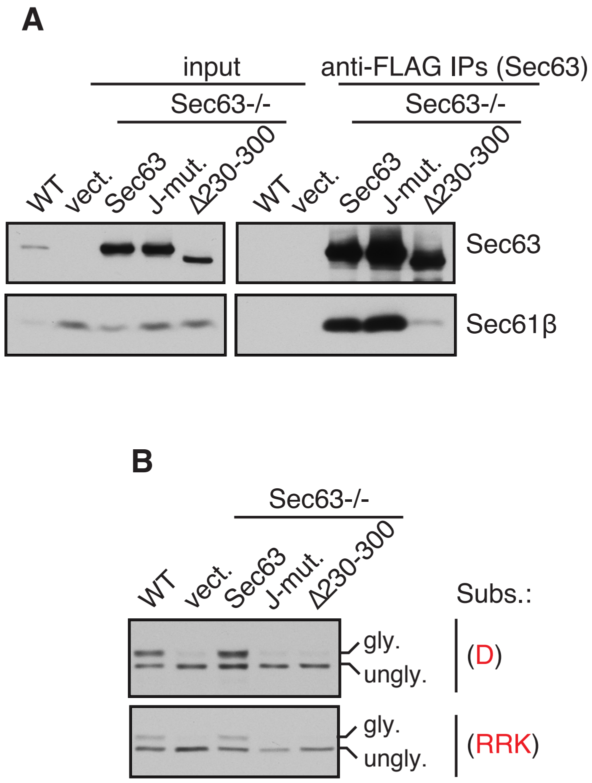


**Figure S2. Both J-domain and translocon interacting region are required for releasing translocon retained substrates.**

(A) WT HEK293 or Sec63-/- cells stably expressing the indicated FLAG-tagged Sec63 constructs were lysed, immuno-precipitated with anti-FLAG beads, and analyzed by immunoblotting for indicated antigens.

(B) The indicated cell lines were transfected with C-TMDs of substrates carrying charged amino acid(s) of either D or RRK. Transfected cells were directly analyzed by immunoblotting with anti-GFP antibodies for substrates.

### Figure S3

**A**

AGAL_HUMAN (100%), 48,768.3 Da
Alpha-galactosidase A OS=Homo sapiens OX=9606 GN=GLA PE=1 SV=1
11 exclusive unique peptides, 14 exclusive unique spectra, 14 total spectra, 152/429 amino acids (35% coverage)

```
MQLRNPELHL GCALALRFLA LVSWDIPGAR ALDNGLARTP TMGWLHWERF
MCNLDCQEEP DSCISEKLFM EMAELMVSEG WKDAGYEYLC IDDCWMAPQR
DSEGRLQADP QRFPHGIRQL ANYVHSKGLK LGIYADVGNK TCAGFPGSFG
YYDIDAQTFA DWGVDLLKFD GCYCDSLENL ADGYKHMSLA LNRTGRSIVY
SCEWPLYMWP FQKPNYTEIR QYCNHWRNFA DIDDSWKSIK SILDWTSFNQ
ERIVDVAGPG GWNDPDMLVI GNFGLSWNQQ VTQMALWAIM AAPLFMSNDL
RHISPQAKAL LQDKDVIAIN QDPLGKQGYQ LRQGDNFEVW ERPLSGLAWA
VAMINRQEIG GPRSYTIAVA SLGKGVACNP ACFITQLLPV KRKLGFYEWT
SRLRSHINPT GTVLLQLENT MQMSLKDLL
```

**B**

CD44_HUMAN (100%), 81,537.5 Da
CD44 antigen OS=Homo sapiens OX=9606 GN=CD44 PE=1 SV=3
6 exclusive unique peptides, 6 exclusive unique spectra, 6 total spectra, 63/742 amino acids (8% coverage)

```
MDKFWWHAAW GLCLVPLSLA QIDLNITCRF AGVFHVEKNG RYSISRTEAA
DLCKAFNSTL PTMAQMEKAL SIGFETCRYG FIEGHVVIPR IHPNSICAAN
NTGVYILTSN TSQYDTYCFN ASAPPEEDCT SVTDLPNAFD GPITITIVNR
DGTRYVQKGE YRTNPEDIYP SNPTDDDVSS GSSSERSSTS GGYIFYTFST
VHPIPDEDSP WITDSTDRIP ATTLMSTSAT ATETATKRQE TWDWFSWLFL
PSESKNHLHT TTQMAGTSSN TISAGWEPNE ENEDERDRHL SFSGSGIDDD
EDFISSTIST TPRAFDHTKQ NQDWTQWNPS HSNPEVLLQT TTRMTDVDRN
GTTAYEGNWN PEAHPPLIHH EHHEEEETPH STSTIQATPS STTEETATQK
EQWFGNRWHE GYRQTPKEDS HSTTGTAAAS AHTSHPMQGR TTPSPEDSSW
TDFFNPISHP MGRGHQAGRR MDMDSSHSIT LQPTANPNTG LVEDLDRTGP
LSMTTQQSNS QSFSTSHEGL EEDKDHPTTS TLTSSNRNDV TGGRRDPNHS
EGSTTLLEGY TSHYPHTKES RTFIPVTSAK TGSFGVTAVT VGDSNSNVNR
SLSGDQDTFH PSGGSHTTHG SESDGHSHGS QEGGANTTSG PIRTPQIPEW
LIILASLLAL ALILAVCIAV NSRRRCGQKK KLVINSGNGA VEDRKPSGLN
GEASKSQEMV HLVNKESSET PDQFMTADET RNLQNVDMKI GV
```

**C**

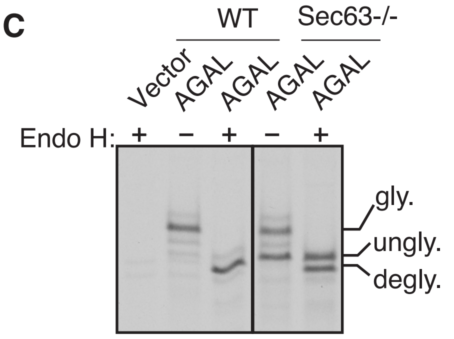


**D**

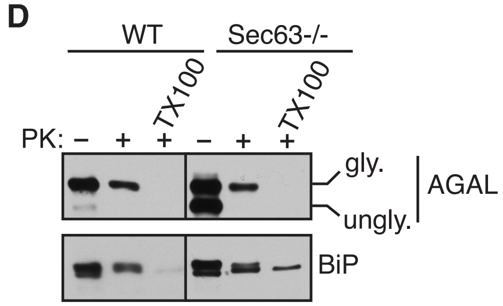


**Figure S3. AGAL mislocalizes to the cytosol in the absence of Sec63**

(A, B) Amino acid sequences of AGAL and CD44 are annotated to indicate the peptides (yellow) identified by mass spectrometry. The signal sequences were underlined.

(C) AGAL expressing cells were immunoprecipitated after radiolabeling for 30 min. The immunoprecipitants were either treated with or without Endo H and analyzed by SDS-PAGE autoradiography. “gly” indicates glycosylated forms. “ungly” denotes non-translocated/unglycosylated AGAL with the uncleaved SS. “degly” indicates Endo H digested de-glycosylated forms.

(D) HEK293 or Sec63-/- cells expressing AGAL were treated with a low concentration of digitonin to selectively permeabilize the plasma membrane followed by incubation with or without proteinase K (PK) and analyzed by immunoblotting. Note that the non-translocated/unglycosylated AGAL with the signal sequence is completely digested by PK because it is localized to the cytosol.

**Figure S4**

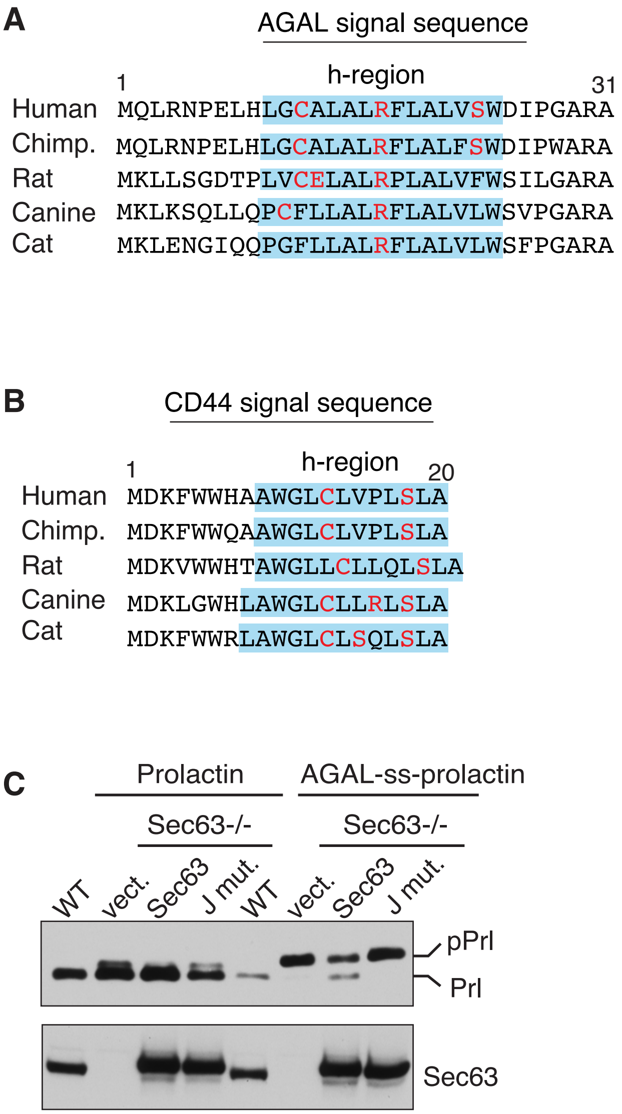


**Figure S4. Conserved weak signal sequences determine Sec63 dependency**
(A, B) Conservation of AGAL or CD44 signal sequence in different mammalian species. Blue shade indicates the hydrophobic region (h-region) of the signal sequence defined by the Kyte-Doolittle scale. Charged and hydrophilic amino acids in the h-region are indicated in red color.
(C) WT HEK293 cells, Sec63-/- cells, or Sec63-/- cells stably expressing wild type Sec63 and or J-domain mutant (HPD/AAA) were transfected with plasmids expressing prolactin or AGAL-ss-prolactin. Cell lysates were analyzed by immunoblotting with an anti-FLAG antibody for substrates or Sec63 antibodies. Precursor (pPrl) and processed (Prl) forms are indicated.

**Figure S5**

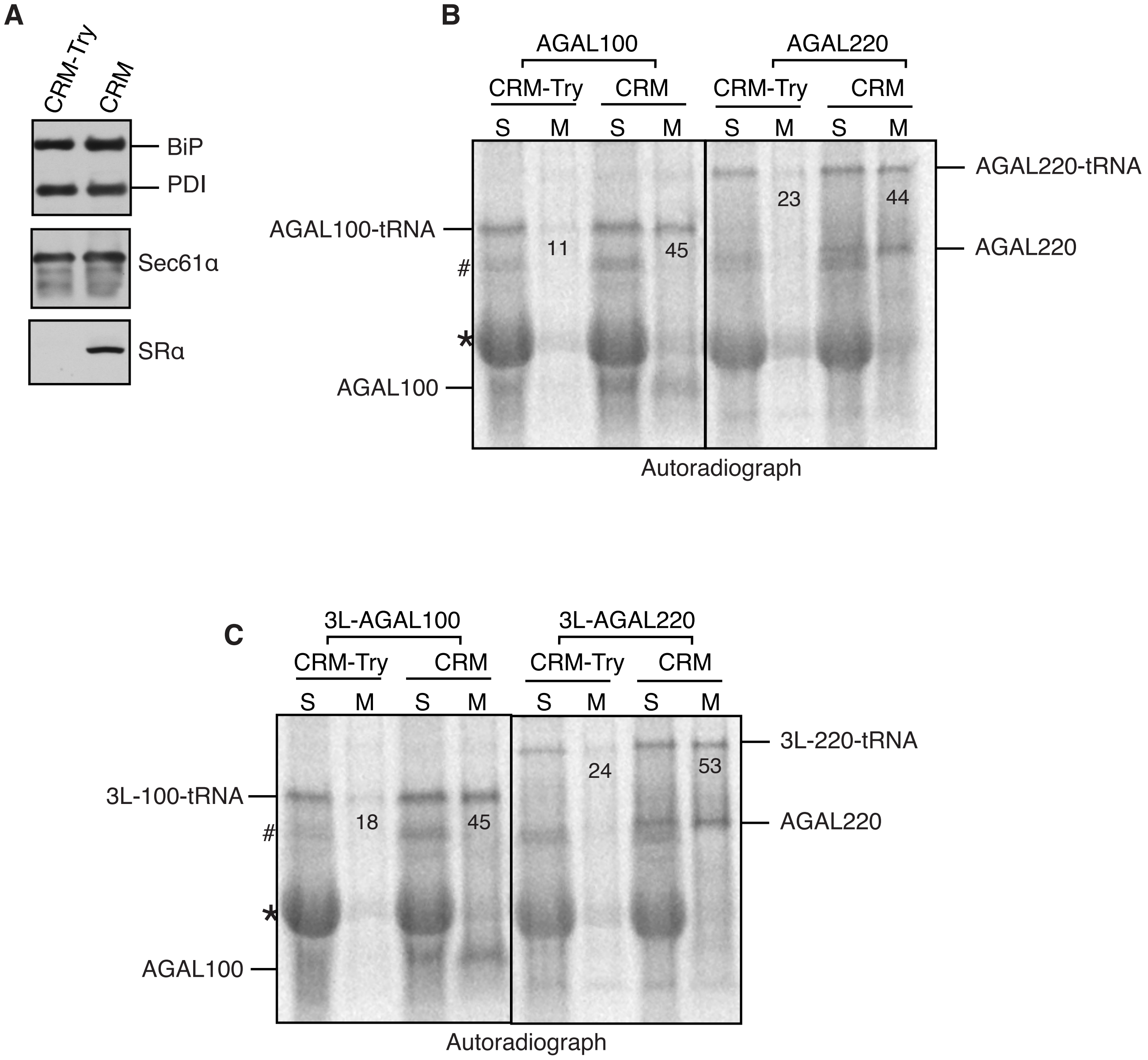


**Figure S5.The weak signal sequence of AGAL can efficiently target its ribosome nascent chain to the ER membrane.**

(A) Untreated and trypsin (20µg/ml) treated canine rough microsomes (CRM) were analyzed by immunoblotting for the indicated antigens. Note that mild trypsin digestion completely shaved off the alpha subunit of the SRP receptor, which is required for the recruitment of the ribosome nascent chains (RNCs) to the ER membrane.

(B, C) RNCs of at indicated lengths of AGAL or 3L-AGAL were produced in RRL in the presence of either trypsin treated CRM (CRM-Try) or untreated CRM. The resulting reactions were layered on 1M sucrose and centrifuged to collect supernatants and pellets, which were then analyzed by SDS-PAGE and autoradiography. The values indicate the percentage of membrane-associated tRNA-linked AGAL or 3L-AGAL. Note that tRNA moieties from AGAL nascent chains could be removed by digestion with RNase A (not shown). Star indicates the background signal from abundant hemoglobulin in RRL. # indicates a non-specific protein.

### Figure S6

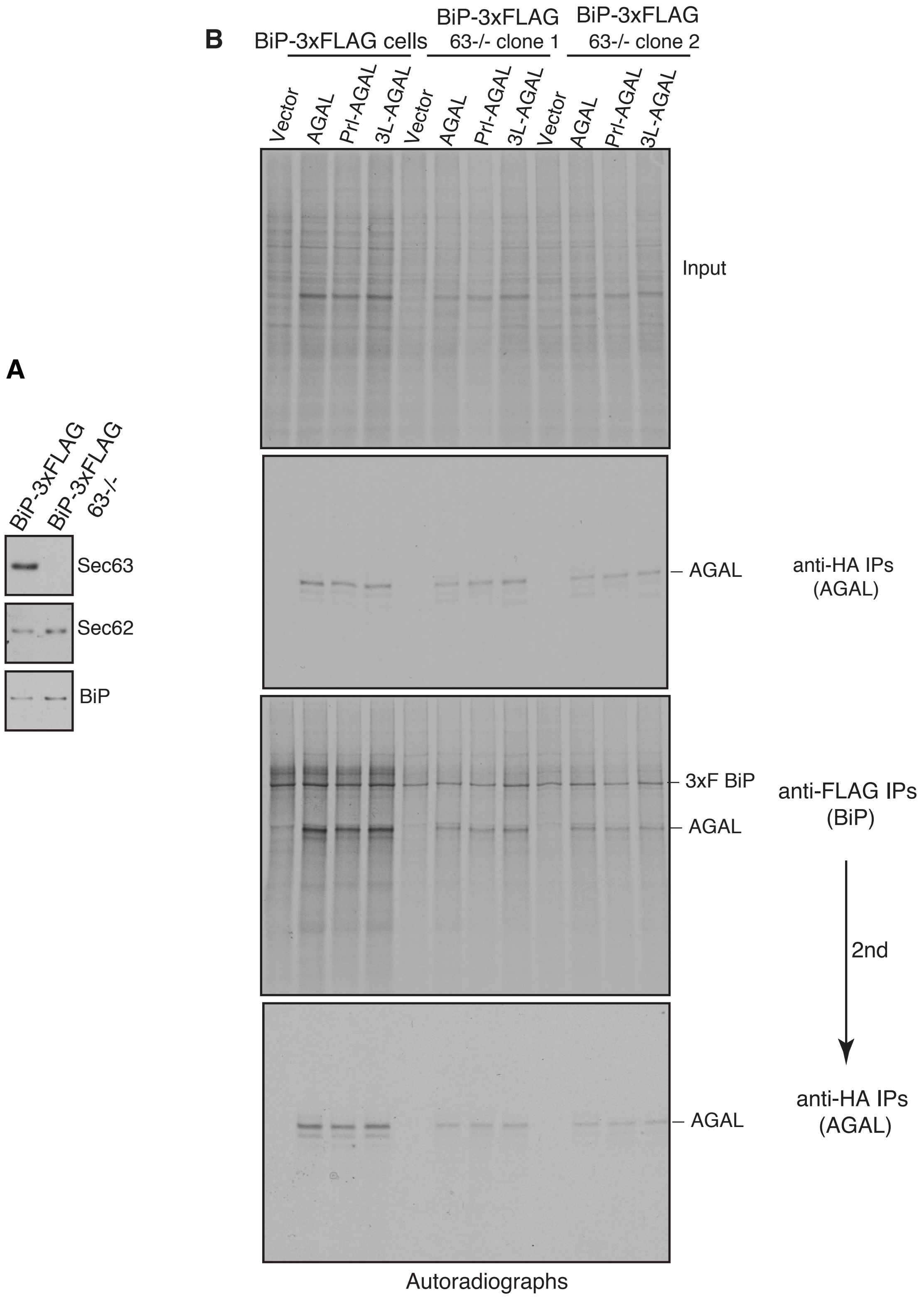


**Figure S6. Sec63 mediates BiP binding to translocating nascent chains containing both weak and strong signal sequences.**

(A) BiP-3xFLAG HEK293 or BiP-3xFLAG HEK293 Sec63-/- cells were analyzed by immunoblotting.

(B) BiP-3xFLAG HEK293 or BiP-3xFLAG HEK293 Sec63-/- cells were transfected with either empty vector or AGAL constructs. Cells were radiolabeled for 2 minutes and immunoprecipitated with either anti-FLAG antibodies for BiP or anti-HA antibodies for AGAL and analyzed by autoradiography. Anti-FLAG immunoprecipitants were diluted and re-immunoprecipitated (2nd IPs) with anti-HA antibodies for AGAL.

### Figure S7

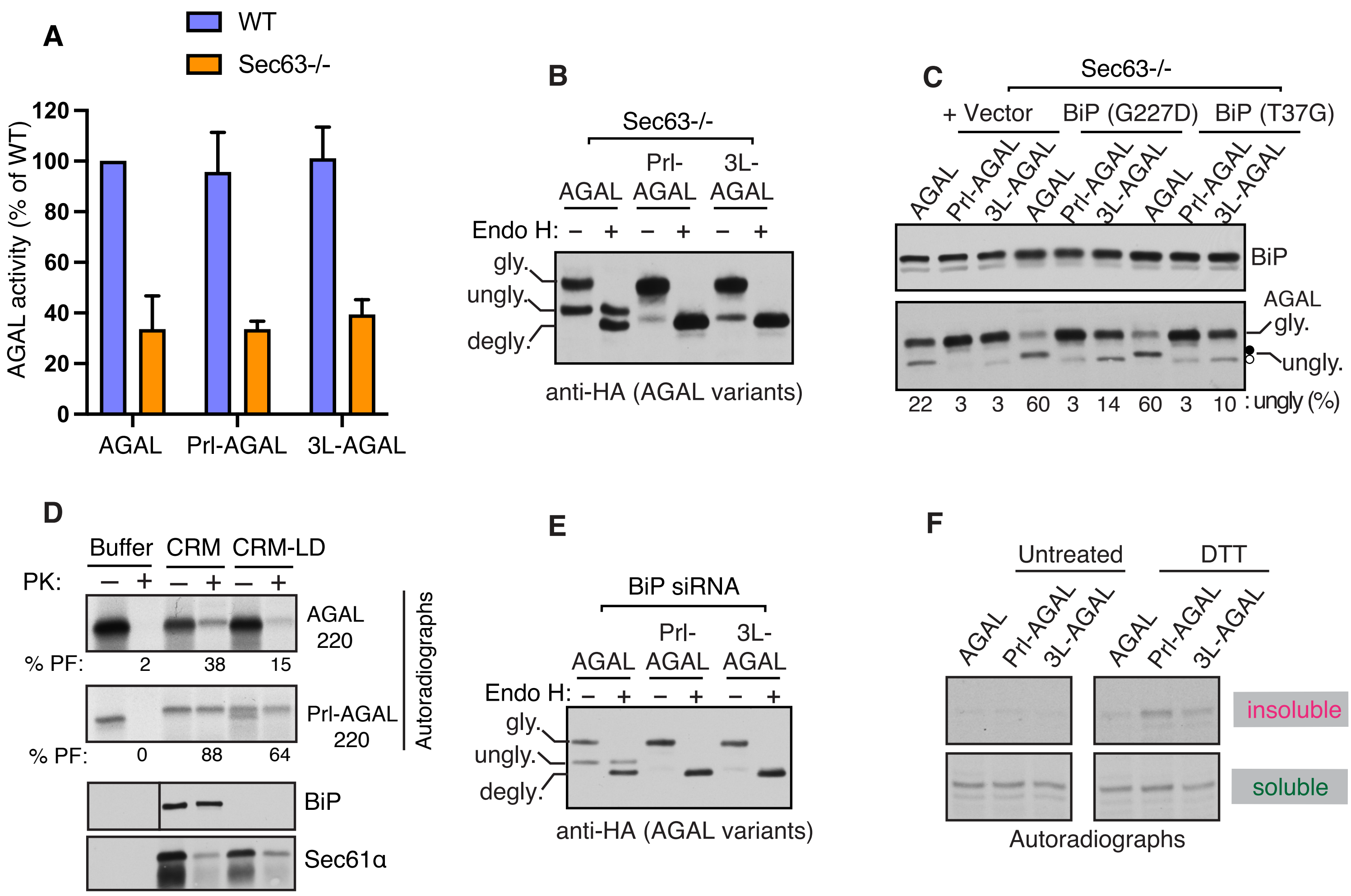


**Figure S7. Effect of signal sequences on protein translocation and protein folding.**
(A) WT or Sec63-/- cells expressing the indicated AGAL constructs were lysed in NP40 buffer and the supernatant (soluble fraction) after centrifugation was assayed for α-galactosidase activity. Data are shown as a percentage of the activity of WT AGAL. Error bar represents the standard deviation (n=3).
(B) Sec63-/- cells expressing the indicated variants of AGAL were treated or not treated with Endo H and analyzed by immunoblotting. "gly" indicates glycosylated forms. "ungly" denotes non-translocated/unglycosylated forms with uncleaved SSs. "degly" indicates Endo H digested deglycosylated forms. Note that the unglycosylated AGAL migrates slower than the Endo H deglycosylated AGAL, suggesting that the unglycosylated form contains the uncleaved SS. However, unglycosylated forms of Prl-AGAL and 3L-AGAL carrying uncleaved SSs migrate only slightly slower than the Endo H digested deglycosylated bands. Presumably, this difference is caused by the presence of a positively charged arginine (R) residue in the middle of the AGAL SS but not in the SS of Prl-AGAL or 3L-AGAL.
(C) Sec63-/- cells co-expressing AGAL constructs along with either vector, BiP (G227D), or BiP (T37G) were analyzed by immunoblotting. The unglycosylated band is given as the percentage of total signals from both glycosylated and unglycosylated bands.
(D) Canine rough microsomes (CRM) were permeabilized with a low concentration of detergent to release the luminal content and were resealed by diluting with a detergent-free buffer and centrifugation. Transcripts encoding AGAL-220 and Prl-AGAL-220 were translated in the presence of indicated membranes or buffer followed by digestion with or without proteinase K (PK) and analyzed by autoradiography. The percentage of protease-protected (PF) chains is shown below each autoradiograph.
(E) Total lysate of BiP depleted HEK293 cells shown in Figure 6E was treated or non-treated with Endo H and analyzed as in B.
(F) Cells expressing the indicated AGAL constructs were treated with or without 10mM DTT, radiolabeled 2min for untreated and 7min for DTT treated cells. Detergent soluble and insoluble fractions were immunoprecipitated with anti-HA antibodies and analyzed by autoradiography.
